# Phosphatase activity is dispensable for PRL-3-mediated oncogenesis and tumor progression

**DOI:** 10.1101/2025.05.14.654016

**Authors:** Jeffery T. Jolly, Majd A. Al-Hamaly, Caroline N. Smith, H. Peter Spielmann, Jessica S. Blackburn

## Abstract

Phosphatase of Regenerating Liver 3 (PRL-3) is frequently upregulated in cancer and associated with poor prognosis, yet its oncogenic mechanism remains unresolved. Although traditionally studied for its phosphatase activity, PRL-3 also engages in protein-protein interactions via its catalytic site, notably binding the CNNM magnesium transporters, rendering these functions mutually exclusive. The commonly used C104S mutant disrupts both activities, confounding interpretations of prior studies. To dissect PRL-3’s distinct functions, we employed mutants selectively deficient in phosphatase activity (C104D) or CNNM binding (R138E). In zebrafish models of acute lymphoblastic leukemia (ALL) and rhabdomyosarcoma (RMS), as well as in human cancer cell lines, both wild-type PRL-3 and C104D enhanced tumor initiation, growth, and dissemination, while R138E had no effect. These findings indicate that PRL-3 promotes cancer via non-catalytic mechanisms. To support therapeutic development, we established an in vitro FRET-based assay to screen for inhibitors of the PRL-3:CNNM interaction and validated a cyclic peptide as a positive control. By demonstrating that PRL-3 enhances cancer progression independently of its catalytic activity, this study shifts focus toward targeting its binding functions as a therapeutic strategy.

**Graphical Abstract:** 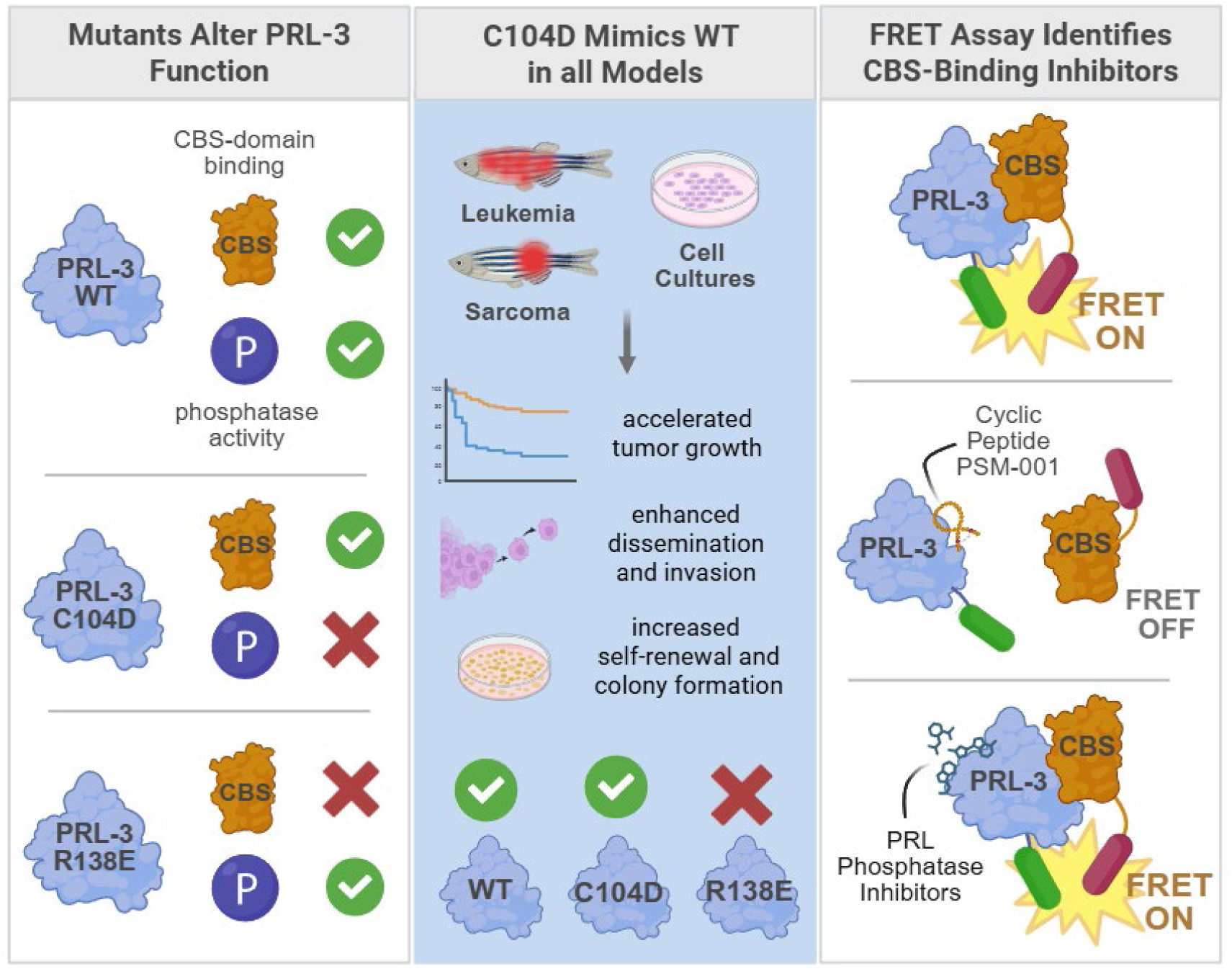

**Statement of significance:** This study demonstrates that PRL-3 drives cancer initiation and progression through a phosphatase-independent mechanism across diverse models, highlighting the need to therapeutically target its non-catalytic protein interactions in future cancer treatment strategies.

## Introduction

Phosphatase of Regenerating Liver 3 (PRL-3) is frequently upregulated in a wide range of cancers and correlates with increased metastasis and poor patient outcomes. Although it is normally silenced in adult tissues, PRL-3 becomes re-expressed in tumors and has been linked to enhanced invasion, dissemination, and therapeutic resistance (1–5). Despite its strong association with cancer progression, the precise molecular mechanism by which PRL-3 promotes malignancy remains unclear (6–11).

Most studies have concentrated on PRL-3’s phosphatase activity and have depended on the C104S mutant, which eliminates catalytic function, to examine its role in cancer progression. Utilizing this mutant, the phosphatase activity of PRL-3 has been implicated in the enhancement of migration, invasion, and colony formation across various models (12–25). Therefore, identifying the cellular substrates of PRL-3 responsible for these effects has been a key area of interest. However, PRL-3 acts on a broad range of substrates, including proteins, lipids, amino acids, and nucleotides, making it difficult to identify a single key effector (26–31).

In addition to its catalytic role, PRL-3 binds with high affinity to the cystathionine-β-synthase (CBS) domain of CNNM magnesium transporters, utilizing residues that overlap with its active site (32–35). This interaction is proposed to reduce CNNM-mediated magnesium efflux while promoting TRPM7-dependent influx (36–42). Importantly, phosphatase activity and CNNM binding are mutually exclusive. When PRL-3 binds to CNNMs, its active site becomes blocked. Upon substrate dephosphorylation, PRL-3 enters a long-lived intermediate state that prevents CNNM interaction (34). Furthermore, recent investigations have found that the widely used C104S mutant disrupts both functions, complicating the determination of which activity drives PRL-3’s cancer-promoting effects (30).

To address this issue, Kozlov et al. characterized PRL-3 mutants that isolate its individual functions in vitro (24). Specifically, they described the PRL-3 C104D mutant, which lacks phosphatase activity but retains CNNM binding, and the R138E mutant, which significantly reduces affinity for binding the CBS domain of CNNMs. Surprisingly, in a B16 melanoma allograft model, PRL-3 C104D promoted metastasis, whereas R138E had no effect. These results suggest that PRL-3 may function, at least in part, through non-catalytic mechanisms. While compelling, the B16 model cannot evaluate early oncogenic events or tumor initiation. Furthermore, the past two decades have implicated the phosphatase activity of PRL-3 in oncogenesis, local invasion, angiogenesis, and resistance to apoptosis across a wide variety of cancer types (43–46). Together, these findings raise the question of whether PRL-3’s phosphatase and non-catalytic functions contribute differently to distinct stages of cancer progression. Determining whether one activity initiates oncogenic events while the other drives dissemination may reveal functionally separable roles for PRL-3 in malignancy. Considerable effort has also focused on developing small molecules targeting PRL-3’s phosphatase activity (47–57); it is critical to determine if inhibiting catalytic function alone is sufficient for therapeutic benefit or whether the protein’s non-catalytic properties must also be addressed.

To determine which function of PRL-3 drives tumor initiation and progression, we used transgenic zebrafish models of acute lymphoblastic leukemia (ALL) and rhabdomyosarcoma (RMS), along with human cancer cell lines. The optical transparency and genetic tractability of zebrafish enabled real-time visualization of early oncogenesis and tumor dissemination in vivo (58–60), while in vitro assays assessed migration, invasion, and anchorage-independent growth. We found that PRL-3 enhanced these cancer-associated phenotypes independently of its phosphatase activity, as effects were maintained in the phosphatase-dead C104D mutant but lost with the CNNM-binding deficient R138E variant. To support future studies, we developed a FRET-based assay using purified proteins to quantify PRL:CNNM binding and validated a cyclic peptide inhibitor. These findings establish a functional separation between PRL-3’s catalytic and protein-binding roles, suggesting that its oncogenic effects are driven by non-catalytic interactions and that disrupting these interactions may be a more effective therapeutic strategy than phosphatase inhibition.

## Materials and Methods

### Publicly Available Data Sets and Web-Based Tools

TCGA PanCancer Atlas data were analyzed via cBioPortal (RRID:SCR_014555) using mRNA z-scores relative to normal samples (log RNA Seq V2 RSEM), with a z-score threshold ≥ 2 (61–63). Overall survival analyses based on PTP4A3 expression were performed using KM-plotter (RRID:SCR_018753), incorporating both RNA-seq and GeneChip datasets (PTP4A3, 206574_s_at) (64–67). Molecular models were generated with AlphaFold3 (RRID:SCR_025885) (68) and adjusted for presentation with PyMOL Molecular Graphics System, Version 3.0 Schrödinger, LLC (RRID:SCR_000305). Protein sequences were obtained from UniProt (RRID:SCR_002380) (69), and aligned with Jalview (RRID:SCR_006459) (70). Graphical illustrations were made with BioRender (RRID:SCR_018361) and Adobe Illustrator (RRID:SCR_010279).

### Plasmids and Cloning

The human PTP4A3 (PRL-3) coding sequence was PCR-amplified and cloned into pENTR-D-TOPO (Thermo Fisher, K240020). Site-directed mutagenesis (Agilent QuikChange Lightning, 210518) was used to generate PRL-3 mutants (see **Supplemental Table 1** for primers). Entry clones were recombined into Rag2.GW.DEST (71), pLPC-N FLAG (RRID:Addgene_12521), pLenti-CMV-Puro-DEST (RRID:Addgene_17452), or pCW57.1 (RRID:Addgene_41393) using LR Clonase II (Thermo Fisher, 11791020). Control mCherry vectors were generated from pME-mCherry (Tol2 Kit) (72). Rag2.m-Myc and Rag2.KRAS.G12D vectors were generated as previously described (73,74). Plasmids were propagated in Top10 *E. coli* (Thermo Fisher, C409601) under kanamycin or ampicillin selection and purified with IBI Miniprep Kit (IBI Scientific, IB47101). All constructs were sequence-verified by Sanger or whole-plasmid sequencing (Eurofins Genomics). A full list of constructs is provided in **Supplemental Table 2**.

### Cell Culture

HEK293T (CRL-3216, RRID:CVCL_0063), RD (CCL-136, RRID:CVCL_1649), HCT 116 (CCL-247, RRID:CVCL_0291), and Jurkat (TIB-152, RRID:CVCL_0367) cell lines were acquired from ATCC and routinely tested for mycoplasma every three months or when a new stable line was established (LookOut® Mycoplasma PCR Detection Kit, Sigma-Aldrich, MP0035-1KT). HEK293T, HCT116, and RD cells were cultured in high-glucose DMEM (Hyclone, SH30022.FS) with 1 mM sodium pyruvate (Thermo Fisher, 11360070) and 10% FBS (Sigma-Aldrich, 12306C) at 37 °C in a humidified 5% CO_2_ environment. Jurkat cells were cultured similarly in RPMI 1640 (Hyclone, SH3002702) with 10% FBS.

### Transfection, Co-immunoprecipitation, and Phosphatase Activity Assay

HEK293T cells (2.5 × 10⁶ per 10 cm dish) were transfected with 5 µg plasmid using Lipofectamine™ 3000 (Thermo Fisher, L3000001) in Opti-MEM (Thermo Fisher, 31985062) and harvested 48 hours later. Cells were lysed in Pierce™ IP Lysis Buffer (Thermo Fisher, 87788) with EDTA and protease inhibitors (Boster, AR1182), incubated on ice for 30 minutes, and precleared by centrifugation (14,000 RCF, 15 minutes). Anti-FLAG® M2 Magnetic Beads (Sigma-Aldrich, M8823, RRID:AB_2637089), blocked with 5% BSA (Sigma-Aldrich, A7030), were incubated with lysates overnight at 4 °C. Beads were washed in PBS and eluted with 4× Laemmli Sample Buffer (Bio-Rad, 1610747) for western blotting.

For phosphatase activity assays, beads were resuspended in reaction buffer (10 mM Tris-HCl, pH 7.5; 15 mM NaCl; 10 mM TECP; 1 mM EDTA) (31) and incubated for one hour at room temperature. DiFMUP (100 µM; EnzChek™ Kit, Thermo Fisher, E12020) was added and incubated for 30 minutes. Supernatants were transferred to black 384-well plates (Corning, 3575) and read at 358/450 nm (ex/em) using a Cytation 5 plate reader (Agilent, RRID:SCR_019732). Fluorescence from DiFMUP-only bead controls was subtracted.

### Nanobody Immunofluorescence Imaging

PRL-3 mutants were LR-cloned into pLenti-CMV-Puro-DEST and transfected into HCT116 cells using Lipofectamine™ 3000, as described above. Localization was assessed via immunofluorescence (IF) microscopy using a PRL-3-specific nanobody (nanobody 91) following established protocols (57). Cells were seeded at 5,000 per well in 96-well black glass-bottom plates (Cellvis, P96-1.5H-N) and fixed 24 hours post-transfection in 4% paraformaldehyde (Thermo Fisher, J61899.AP) for 15 minutes, followed by PBS washes. Permeabilization was performed with 1% Triton X-100 (Sigma, X100-100) for 10 minutes. Cells were blocked in 2% BSA/PBS for one hour before incubation with nanobody 91 (1 mg/mL stock, diluted 1:100 in blocking buffer) for one hour at room temperature. After washing with PBS, cells were labeled with Alexa Fluor-594–conjugated anti-alpaca IgG VHH (Jackson ImmunoResearch, 128-585-232; 1:400) and counterstained with Hoechst 33342 (Thermo Fisher, H3570; 1:1000). After five PBS washes, imaging was performed on a Nikon A1R confocal microscope with a 40× water objective at the University of Kentucky Light Microscopy Core (RRID:SCR_026405). Image adjustments (brightness only) were applied uniformly with Fiji (ImageJ, v1.54, RRID:SCR_002285) Channels were assigned as follows: Hoechst (405 nm), GFP-PRL-3 (488 nm), and nanobody (561 nm).

### Zebrafish Husbandry and Microinjection for RMS and ALL Models

Zebrafish were maintained on a 14:10 light/dark cycle at 28 °C under University of Kentucky IACUC approval (Protocol 2019–3399) (RRID:NCBITaxon_7955). The Casper strain (ZDB-FISH-150901-6638) was used for all cancer models (75). In addition, p53 knockout Casper animals were used as a positive control for enhanced tumorigenesis in the RMS model (76). Rag2 gateway vectors encoding mCherry or PRL-3 mutants were linearized with XhoI (NEB, R0146) and purified by phenol/chloroform extraction. Injection mixes (0.5 nL) contained 40 ng/µL each of Rag2:mCherry, Rag2:gene of interest (PRL-3 variant or additional mCherry for the control group), and either Rag2:m-Myc (ALL model) or Rag2:h-KRAS.G12D (RMS model) in injection buffer (5 mM Tris-HCl, 0.5 mM EDTA, 100 mM KCl). Single-cell Casper embryos were injected within 30 minutes post-fertilization using a Narishige microinjection system.

### Live Animal Imaging and Analysis

Animals were assessed for mCherry-positive cells at 15 dpf using a Nikon SMZ25 fluorescence microscope and imaged every five days while anesthetized with 0.015% MS-222 (tricaine methanesulfonate) (TCI, T0941). Circulating ALL (penetrance) is defined as detecting mCherry-positive cells circulating in the tail fin vasculature within a 5-second observation frame. ALL animals were monitored for up to 90 days. Animals were euthanized upon reaching >60% leukemia burden. RMS animals were monitored for up to 60 dpf (no new mCherry-positive animals arose after 15 dpf). Outliers in either group with >75% mCherry-positive bodies at 15 dpf were excluded from the analysis and euthanized. Fiji (ImageJ, v1.54) was used to quantify thymus size (ALL) as well as tumor size and morphology (RMS) (77).

### RNA Extraction and Quantitative PCR

Animals were euthanized with 0.05% MS-222 before cancer cell isolation. ALL cells were collected by mincing tissue (post-head removal), filtering through a 40 μm strainer (Fisher Scientific, 22363547), and pelleting in PBS + 20% FBS. Wild-type whole blood (WB) was prepared similarly by pooling three age-matched Casper animals. RMS tumors and control muscle tissue were dissected, flash-frozen, and stored at –80 °C. RNA was extracted using the Zymo Quick-RNA Miniprep Kit (Zymo, R1054) following genomic DNA removal, and quantified with the Qubit™ RNA HS kit (Thermo Fisher, Q32852). cDNA was synthesized using the iScript™ cDNA Synthesis Kit (Bio-Rad, 1708890). qPCR was performed with 12.5 ng cDNA, 0.25 μM primers (**Supplemental Table 3**), and Forget-Me-Not™ EvaGreen® qPCR Master Mix (Biotium, 31043) on a Bio-Rad CFX384 Touch Real-Time PCR System (RRID:SCR_018057) (40 cycles, SYBR channel). Cq values were normalized to zebrafish housekeeping genes (*z-eef1a* and *z-rplp0*). Fold change was calculated as:

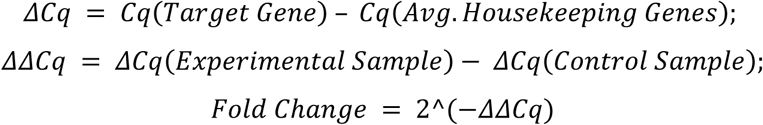

### May-Grünwald Giemsa Stain for Leukemia Samples

Leukemic animals with 40–60% burden were euthanized with 0.05% MS-222, and cells were extracted as described above. Cells were resuspended at 5×10⁵ cells/mL in RPMI and loaded into EZ Single Cytofunnels (Epredia, A78710003) with coated cytoslides (Thermo, 3319). Samples were centrifuged at 1000 rpm for 5 minutes using a Thermo Shandon Cytospin 3 (RRID:SCR_018628). Staining was performed following standard procedures (78). Briefly, Slides were stained with May-Grünwald for 5 minutes, rinsed, and stained with Giemsa for 20 minutes before air-drying overnight. Images were acquired using a Lionheart FX microscope (RRID:SCR_019744).

### Histology and H&E Staining

RMS tumor-bearing animals were sacrificed at 45–55 dpf using 0.05% MS-222. Sample processing was performed by the University of Kentucky Pathology Research Core (RRID: SCR_018824). Whole animals were fixed in 10% neutral buffered formalin (MilliporeSigma, HT501128-4L) for 24 hours at 4 °C, then transferred to 70% ethanol and processed with an Excelsior AS tissue processor (Epredia). Samples were embedded longitudinally in paraffin, sectioned at 4 µm using a manual microtome, and mounted onto positively charged slides. Slides were dried upright at 58 °C for ≥ one hour and stained with hematoxylin and eosin using a Tissue-Tek Prisma stainer. Images were acquired with a Lionheart FX microscope.

### Annexin V and EdU FACS-Based Assays

ALL-bearing animals were euthanized at 30–35 dpf using 0.05% MS-222. Cells were extracted by mincing and filtering tissues through a 40 µm strainer, then maintained in RPMI 1640 with 20% FBS at room temperature unless otherwise noted. For viability analysis, cells were incubated with 0.1 µg/mL DAPI (Sigma-Aldrich, D9542) for 15 minutes at room temperature, rinsed with PBS and annexin-binding buffer (Invitrogen, V13242), and resuspended at 2.5 × 10⁶ cells/mL on ice. Annexin V FITC (TONBO, 35-6409-T100) was added at 0.1 µg/100 µL, incubated 15 minutes on ice in the dark, and diluted with binding buffer for immediate analysis. For EdU labeling, cells were treated with DAPI as above, rinsed with PBS, and resuspended at 2.5 × 10⁶ cells/mL in culture media. EdU (10 µM) was added for one hour at room temperature. Cells were then fixed, permeabilized, and stained using the Click-iT Plus EdU Alexa Fluor 488 Flow Cytometry Kit (Invitrogen, C10632), per manufacturer’s instructions. FACS was performed on a BD FACSymphony A3 (RRID:SCR_023644), and data were analyzed with FlowJo (RRID:SCR_008520). Tumor cells were identified by mCherry positivity, and DAPI-positive dead cells were excluded from analysis.

### Lentivirus and Inducible Cell Line Generation

The pCW57.1 constructs were co-transfected with psPAX2 (RRID:Addgene_12260) and pCMV-VSV-G (RRID:Addgene_8454) into HEK293T cells using Lipofectamine™ 3000. Filtered (0.45 µm) viral supernatants were added to RD and Jurkat cells with 5 µg/mL polybrene (MilliporeSigma, TR-1003-G) and spin-infected (1000 RCF, 1.5 hr). After 48 hours, cells were selected in 1 µg/mL puromycin (Invitrogen, ant-pr-1) for at least 10 days and maintained in puromycin thereafter. Expression was induced with 1 µg/mL doxycycline hyclate (Sigma-Aldrich, D5207-1G) for ≥24 hours. Induction was verified via mCherry fluorescence (EVOS® FL Auto, Life Technologies) or western blotting. All lentiviral procedures were conducted under BSL-2 conditions.

### Western Blotting and Antibodies

Cells were lysed in M-PER (Thermo Fisher, 78503) for standard western blotting or in Pierce IP lysis buffer (as above) with EDTA and protease inhibitors (Boster, AR1182) for co-IP samples. Lysates (20–50 µg) or 5 µL co-IP eluates were loaded onto 4–20% Mini-PROTEAN® TGX Stain-Free™ gels (Bio-Rad, 4568095) and run in Tris/glycine/SDS buffer (Bio-Rad, 1610771EDU). Proteins were transferred to PVDF membranes (Bio-Rad, 1620255) using a Trans-Blot® Turbo™ system (Bio-Rad, 1704150EDU)( RRID:SCR_023156). Blots were blocked with 5% milk in 0.1% TBST (RPI, T32020) for one hour and incubated overnight at 4 °C with primary antibodies: anti-DYKDDDDK (Flag-Tag epitope) (Proteintech, 20543-1-AP; 1:10,000) (RRID:AB_11232216), anti-CNNM3 (Proteintech, 13976-1-AP; 1:1000) (RRID:AB_2082153), anti-α-tubulin (Proteintech, 66031-1-Ig; 1:10,000) (RRID:AB_11042766), or anti-PRL-3 (R&D Systems, MAB3219; 1:1000) (RRID:AB_2174662). After TBST washes, blots were incubated with HRP-conjugated secondary antibodies (anti-mouse IgG, Cell Signaling, 7076P2; anti-rabbit IgG, GE Healthcare, NA9340V) (RRID:AB_330924 and RRID:AB_772191) at 1:2000 in 5% milk for one hour. Blots were developed with Clarity Western ECL Substrate (Bio-Rad, 1705060) and imaged using a ChemiDoc™ Touch Imaging System (Bio-Rad) (RRID:SCR_021693). Chemiluminescent and colorimetric ladder overlays (Bio-Rad, 1610373) were used to generate composite images.

### In Vitro Cell Proliferation Assays

Stable Jurkat, RD, and HCT116 cells were pretreated with or without 1 µg/mL doxycycline hyclate for 24 hours to induce expression. Cells were seeded at 10,000 per well in 48-well plates and incubated in 5% CO₂ for the duration of the assay. Nuclei were stained with Hoechst 33342 (Thermo Fisher, H3570; 4 µg/mL, 20 min) for imaging and cell counting. Jurkat cells were transferred to poly-D-lysine–coated plates (Thermo Fisher, A3890401) and centrifuged (750 g, 10 min) prior to imaging. RD and HCT116 cells were washed with PBS and imaged in the original wells. Cells were imaged using a BioTek Lionheart FX microscope (4× objective, 3×3 montage, DAPI filter), and nuclei were counted with Fiji (ImageJ).

### MethoCult Colony Formation Assay

Jurkat cells (1,000 in 1.25 mL) were suspended in MethoCult supplemented with 50% RPMI, 20% FBS, and 1 µg/mL doxycycline. Cell suspensions (1.25 mL/well) were plated in triplicate 6-well plates; void spaces were filled with sterile water to prevent evaporation. Plates were incubated for 2 weeks in a humidified 5% CO₂ environment. Colonies were imaged on a BioTek Lionheart FX microscope using a 2.5× objective (6 fields/well, montage). Colony number and size were quantified in Fiji (ImageJ), with only colonies ≥50 µm in diameter included in analysis.

### Anchorage-Independent Growth Assay

Soft agar assays were performed in triplicate 6-well plates. Wells were coated with 500 µL of 0.8% agarose (Thermo Fisher, BP160-500) in DMEM and allowed to solidify. RD (50,000) or HCT116 (25,000) cells were suspended in 1.25 mL of 0.4% agarose in DMEM (+10% FBS, 1 mM sodium pyruvate) with 1 µg/mL doxycycline, layered on top, and allowed to set. Void spaces were filled with sterile water to minimize evaporation.

Plates were incubated in 5% CO₂ for 2 weeks (HCT116) or 3 weeks (RD). Fresh media with 1 µg/mL doxycycline was added every 5 days. Colonies were imaged using a BioTek/Agilent Lionheart FX microscope. Colony number and size were quantified in Fiji (ImageJ), with a 50 µm diameter cutoff for inclusion.

### Jurkat Migration Assay

Jurkat cells were cultured in serum-free RPMI with 1 µg/mL doxycycline for 24 hours to induce expression. For the assay, 100,000 cells were seeded in serum-free RPMI with 1 µg/mL doxycycline into the upper chamber of 12-well transwell inserts (0.33 cm², 3 µm pores; Corning, 3472). Lower chambers contained 1 mL of RPMI with 20% FBS and 20 ng/mL CXCL12 (Sino Biological, 10118-HNAE) as chemoattractant. After 16 hours of migration, Hoechst 33342 (4 µg/mL) was added to the bottom chamber for 20 minutes to stain nuclei. Contents were mixed to detach adherent cells and transferred to poly-D-lysine–coated 24-well plates, followed by centrifugation (750 g, 10 min). Cells were imaged in a single plane, and migrated cell numbers were normalized to the mCherry control group.

### RD and HCT116 Invasion Assay

RD and HCT116 cells were cultured in serum-free media with 1 µg/mL doxycycline for 24 hours before the assay. Transwell inserts (8.0 µm pores; Corning, 3464) were coated with 20 µL of 3 mg/mL growth factor-reduced Matrigel (Corning, 47743-718) and incubated for one hour. An additional 5 µL Matrigel was added to fill the meniscus and allowed to solidify for another hour to create a uniform surface. RD (100,000) and HCT116 (150,000) cells were seeded in serum-free DMEM with 1 µg/mL doxycycline into the upper chambers. Bottom wells contained DMEM with 20% FBS as a chemoattractant. Cells were allowed to invade through the matrix for 48 hours. After incubation, non-invading cells and Matrigel were removed from the upper side of the insert with cotton swabs. Invaded cells on the bottom of the insert were stained with Hoechst 33342 and imaged. Cell counts were normalized to the mCherry control for each experiment.

### Recombinant Protein Vectors

The pSKB3 plasmid, a modified pET-28b vector containing a TEV protease site in place of the thrombin cleavage site, was used for recombinant protein production and was a gift from Dr. Konstantin Korotkov (79). Full-length human PRL-3 or HA-CBS was cloned into pSKB3 via NheI/XhoI and T4 ligase. HA-CBS encodes the CBS domain of CNNM3 (residues Y301–D451; RefSeq NM_017623.5) with an N-terminal HA tag and was synthesized as a gBlock (IDT). LSSmGFP-PRL-3.C104D includes large-stokes shift GFP (LSSmGFP) (80) fused to the N-terminus of PRL-3 C104D via a glycine/serine linker. The C104D variant, which lacks an active site cysteine, was used to avoid potential oxidation artifacts. HA-CBS-mScarlet3 includes the HA-CBS domain fused to mScarlet3 (81) at the C-terminus, also joined by a glycine/serine linker. Both fusion constructs were codon-optimized for *E. coli* (IDT Codon Optimization Tool, https://www.idtdna.com/pages/tools) and synthesized as gBlocks with NheI/XhoI sites for cloning. Sequences are provided in **Supplemental Table 2**.

### Generation and Purification of Recombinant Proteins

pSKB3 expression plasmids were transformed into One Shot BL21 Star DE3 *E. coli* (Invitrogen, C601003) and cultured in Lennox LB medium (Millipore Sigma, L3022) with appropriate antibiotics at 37 °C. Cultures were induced at OD₆₀₀ ≈ 0.6 with 0.5 mM IPTG (Fisher, BP175510) and incubated at 16 °C for 16 hours. Cells were harvested by centrifugation (10,000 RCF, 4 °C; Sorvall LYNX 4000, Thermo, 75006580) and resuspended in lysis buffer (300 mM NaCl [VWR, BDH9286], 20 mM Tris pH 7.5, 10 mM imidazole pH 8.0 [Sigma, I2399], 1:1000 protease inhibitor cocktail [Sigma, P8465], 10% glycerol [MilliporeSigma, 0854-1L]). Cells were lysed twice via sonication (Sonics, VCX 750) on ice. Lysates were cleared by centrifugation (38,000 RCF, 4 °C, 50 min), and supernatants were loaded onto Ni-NTA resin (VWR, 786-940) in Econo-Pac Columns (Bio-Rad, 7321010). Proteins were eluted using a buffer with 250 mM imidazole. His-tags were cleaved with TEV protease, and samples were dialyzed overnight at 4 °C in SnakeSkin tubing (Thermo, 68100) against imidazole-free buffer. A second Ni-NTA purification removed uncleaved and contaminating proteins. Final purification was performed on a Superdex 200 Increase 10/300 GL column (GE, 28990944) using an ÄKTA system (GE Healthcare, RRID:SCR_019958) in 100 mM NaCl, 20 mM HEPES pH 7.5 (Fisher, BP310-100). Protein fractions were assessed using 4–20% Mini-PROTEAN TGX Stain-Free Gels (Bio-Rad, 4568094) on a ChemiDoc Touch system (Bio-Rad, 1708370), visualized by Stain-Free or Coomassie staining (MilliporeSigma, B0770; Thermo, 424040025). The purest fractions were pooled, flash-frozen in liquid nitrogen, and stored at −80 °C.

### Cyclic Peptides-PRL Substrate Mimics (PSM): Design and CD Analysis

Cyclic peptides were designed based on the PRL:CNNM interaction interface from the crystal structure (30), incorporating the conserved CNNM3 loop region (residues V416–L432) with variable linker strategies (see **Supplemental Table 4**). Peptides were synthesized by LifeTein (≥98.02% purity). Peptides were dissolved in 10 mM ammonium bicarbonate, then diluted to 0.4 mg/mL in 1 mM NaCl and 200 µM HEPES pH 7.5. Samples (400 µL) were loaded into 1 nm path length quartz cuvettes (Strana Cell, 21-I-1) and analyzed using a Jasco J-810 Circular Dichroism Spectropolarimeter. Far-UV spectra (180–260 nm) were collected at 1 nm intervals at 25 °C. Spectra were normalized to buffer-only controls and averaged from five technical replicates.

### In Vitro FRET Assay: Generation and Validation

LSSmGFP-PRL3.C104D and HA-CBS-mScarlet3 were resuspended in buffer (100 mM NaCl, 20 mM HEPES pH 7.5, 5% glycerol). LSSmGFP-PRL3.C104D (250 nM) was preincubated with test compounds for 30 minutes at room temperature, followed by the addition of HA-CBS-mScarlet3 (500 nM). After 30 minutes, samples were transferred to black 384-well plates (Corning, 3575). FRET signal was measured using a Cytation 7 (BioTek/Agilent) at 400/600 nm (Ex/Em). Wells containing LSSmGFP-PRL3.C104D alone were used for background subtraction. RFU values were normalized to control wells containing DMSO-treated LSSmGFP-PRL3.C104D + HA-CBS-mScarlet3.

### In Vitro Phosphatase Assay

Black 384-well plates were loaded with 0.5 µM PRL-3 in buffer (10 mM Tris-HCl pH 7.5, 15 mM NaCl, 0.01% Triton-X, 5 mM TCEP) and supplemented with test compounds, as previously described (31). Reactions were initiated with 25 µM DiFMUP and fluorescence was measured at 360/450 nm (Ex/Em) for one hour using a Synergy LX Multi-Mode Reader (BioTek/Agilent) (RRID:SCR_019763). Raw RFU values were converted to product concentration (nM) using a DiFMU standard curve from the EnzChek Phosphatase Assay Kit. Experiments were performed twice with independently purified PRL-3 and pooled for analysis. Compounds tested included HA-CBS, cyclic peptides, JMS-053 (MedChemExpress, HY-135457), and Candesartan Cilexetil (Selleckchem, S2037).

### In Vitro Binding Assays

Purified HA-CBS or HA-CBS-mScarlet3 (2.5 µM) was incubated for one hour at room temperature with BSA-blocked HA-Tag (C29F4) magnetic beads (Cell Signaling, 11846S, Lot #9) (RRID:AB_2665471) in IP buffer (25 mM Tris-HCl pH 7.4, 150 mM NaCl, 1% NP-40 [Abcam, ab142227], 5% glycerol [VWR, 0854]). Equimolar PRL-3 or LSSmGFP-PRL3.C104D was prepared in IP buffer with 1 mM TCEP and pre-incubated with 50 µM test compounds for 30 minutes before being added to the HA-CBS-loaded beads for an additional one hour with rotation. Immuno-complexes were captured using a magnetic rack, washed three times, and eluted in 50 µL 4× Laemmli buffer with 2-mercaptoethanol, followed by boiling at 95 °C for 5 minutes. As a negative control, Protein A magnetic beads (Cell Signaling, 73778S) were pre-loaded with rabbit IgG (Cell Signaling, 3900S, Lot #37) (RRID:AB_1550038) and incubated with PRL-3 under identical conditions to assess nonspecific binding. All eluates were analyzed by SDS-PAGE and Western blot as described above. Band intensities were quantified using Fiji (ImageJ) and normalized to DMSO-treated controls.

### Statistical Analysis and Reproducibility

Graphing and statistical analysis were performed using GraphPad Prism (v10.4.1) (RRID:SCR_002798) as detailed in the figure legends. Analyses included Student’s t-test, one- and two-way ANOVA, and Kaplan-Meier survival analysis with two-tailed analysis and assumed normality where applicable. All error bars represent standard deviation. Statistical significance was defined as (*, *p*<0.05; **, *p*<0.01; ***, *p*< 0.001; ****, p< 0.0001). Experimental replicates per analysis were performed as described in the figure legends.

## Results

### PRL-3 is the most frequently upregulated PRL/CNNM family member across cancers and correlates with poor patient prognosis

To justify our mechanistic focus on PRL-3, we first evaluated the relative expression of PRL family members and their CNNM binding partners across a variety of cancers. Using TCGA PanCancer Atlas data accessed via cBioPortal, we assessed mRNA expression (z-score ≥2) in tumor samples relative to normal tissue for all PRLs (PTP4A1–3) and CNNMs (CNNM1–4). Among these genes, PRL-3 (PTP4A3) was the most frequently overexpressed across cancer types, including colorectal, gastric, bladder, and breast cancers (**Figure 1A**), consistent with previous studies linking PRL-3 to cancer progression. Despite its frequent upregulation, PRL-3 showed low rates of genomic amplification and rare mutation across TCGA cohorts (**Figure 1B–C**), suggesting transcriptional regulation likely underlies its increased expression. We next assessed whether PRL-3 overexpression correlates with aggressive tumor features. PRL-3 expression was significantly elevated in tumors with distant metastasis (M1 vs. M0; *p*<0.0001, **Figure 1D**) and increased with histologic tumor grade (G1–G4; *p*<0.0002, **Figure 1E**). Furthermore, high PRL-3 expression was associated with reduced overall survival in multiple cancer types, with hazard ratios ranging from 1.3 to 2.76 (**Supplemental Figure 1**). Together, these data reinforce PRL-3 as a clinically relevant oncogene with consistently elevated expression across multiple cancer types. PRL-3 expression correlates with metastatic potential, higher tumor grade, and poor patient prognosis, reinforcing the need to clarify its molecular mechanism of action.

**Figure 1.**
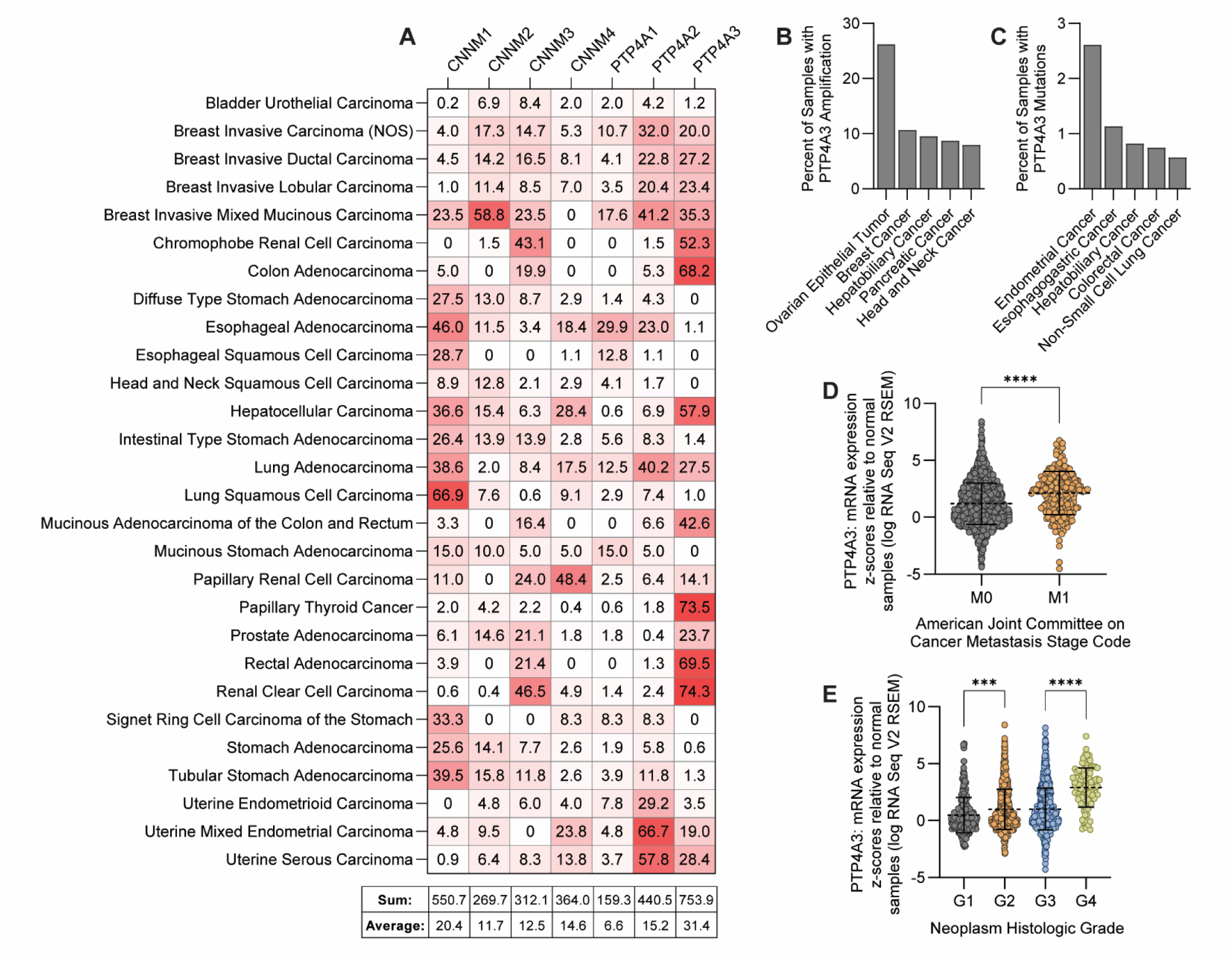
PRL-3 is frequently upregulated in a variety of cancers and correlates with indicators of tumor aggressiveness. **(A)** Heat map showing the percentage of tumor samples within each TCGA cancer type (rows) exhibiting >2-fold mRNA overexpression of CNNM (CNNM1–4) and PRL (PTP4A1–3) family genes (columns), relative to matched normal tissue. Summary statistics for each gene are provided below the heat map. **(B-C)** Frequency of PRL-3/PTP4A3 genomic amplifications (B) and mutations (C) across TCGA cancer types (top 5 shown for each). **(D)** PRL-3/PTP4A3 mRNA expression (z-score relative to normal tissue) in tumors with (M1) or without (M0) distant metastasis, based on American Joint Committee on Cancer staging.**(E)** PRL-3/PTP4A3 mRNA expression across histologic tumor grades (G1-G4; G1 = well differentiated, G4 = undifferentiated). Each data point represents an individual patient sample. Error bars indicate standard deviation. Statistical significance was assessed one-way ANOVA with Tukey’s correction (***, *p*<0.001; ****, *p*<0.0001).

### PRL-3 mutations can distinguish between phosphatase and CNNM binding functions within the cell

To dissect cellular functions of PRL-3, we validated a panel of point mutants previously characterized in vitro for their effects on phosphatase activity and binding to the CBS domain of CNNM (24,82) **Figure 2A-B** summarize these mutations and their positions relative to the CNNM binding interface. While prior studies primarily used purified recombinant proteins under controlled conditions, it remains unclear whether these mutants retain their functional specificity in cells or if mutations can be combined without compromising function.

**Figure 2.**
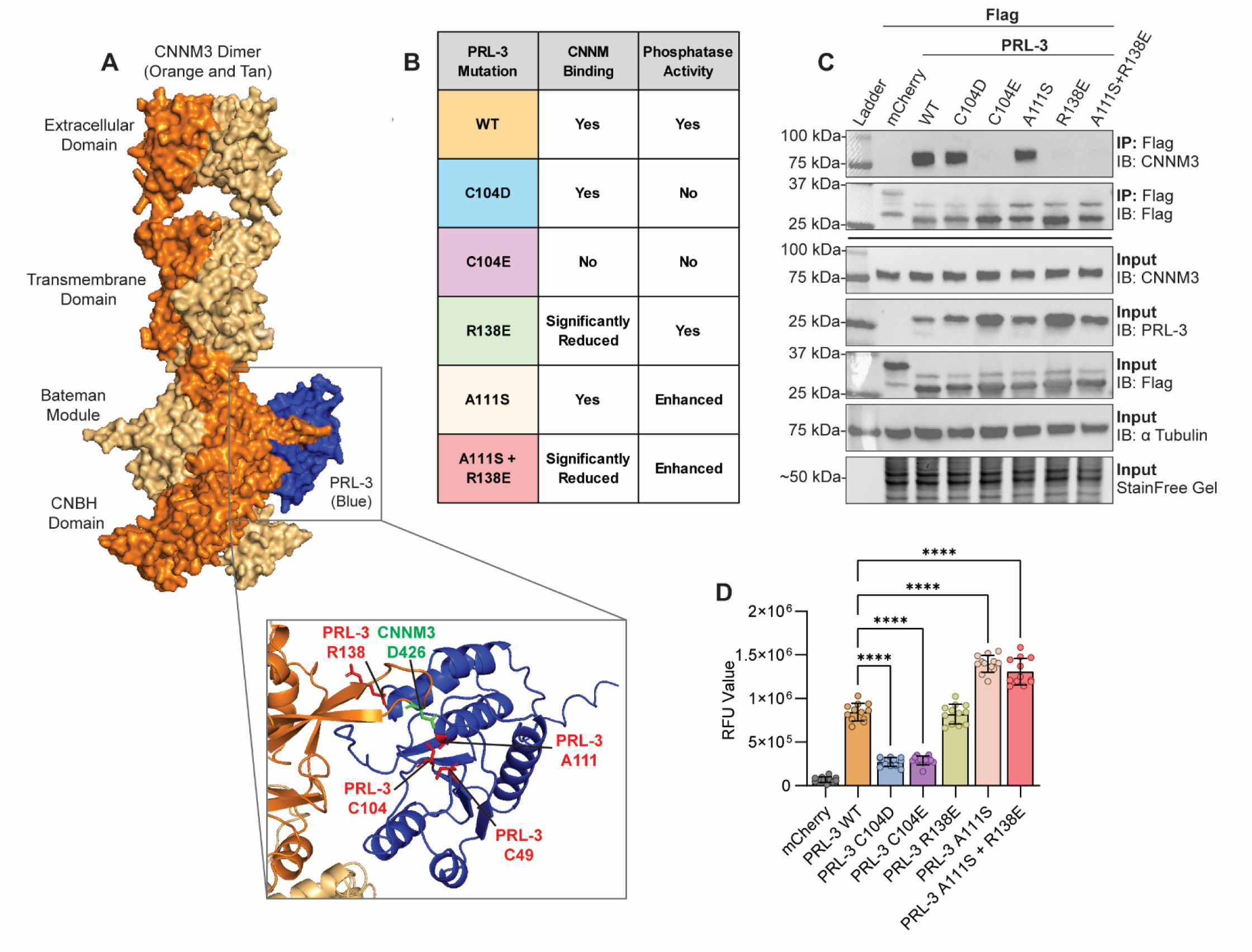
Point mutations delineate PRL-3’s phosphatase activity and CNNM binding functions. **(A)** Structural model of the PRL:CNNM complex rendered with AlphaFold3. CNNM3 homodimers are shown in shades of orange; PRL-3 is in blue. Key interface residues are highlighted in green (CNNM3) and red (PRL-3). **(B)** Summary table of PRL-3 point mutations and their known effects on catalytic activity and CNNM binding. **(C)** Co-immunoprecipitation of FLAG-tagged PRL-3 constructs expressed in HEK293T cells, followed by immunoblotting for endogenous CNNM3. The images shown are representative of at least two independent experiments. **(D)** Phosphatase activity of immunoprecipitated FLAG-tagged PRL-3 mutants, assessed using the fluorogenic substrate DiFMUP (6,8-difluoro-4-methylumbelliferyl phosphate). Data are the pooled results of two biological replicates, with each data point representing the average RFU value of three replicate wells. Error bars indicate standard deviation. Statistical significance was determined by one-way ANOVA with Tukey post hoc correction. ****, *p*<0.0001.

To test this, we transfected HEK293T cells with FLAG-tagged PRL-3 constructs, followed by immunoprecipitation of the constructs to assess endogenous CNNM binding and phosphatase activity. Co-immunoprecipitation revealed that PRL-3 WT, C104D (phosphatase dead, CNNM-binding competent), and A111S (enhanced phosphatase) bound to endogenous CNNM3, whereas C104E (double-deficient), R138E (phosphatase active, CNNM-binding deficient), and A111S+R138E (enhanced phosphatase and reduced CNNM binding) did not (**Figure 2C**). Phosphatase activity assays using the synthetic substrate DiFMUP confirmed that PRL-3 WT and R138E retained catalytic activity, while C104D and C104E mutants lacked activity (*p*<0.0001). A111S exhibited significantly higher phosphatase activity than WT, an effect maintained in the A111S+R138E double mutant (**Figure 2D**, *p*<0.0001). These data confirm that these PRL-3 mutants retain their expected functional properties in a cellular context, supporting their use as tools to dissect PRL-3’s distinct activities in vivo.

To ensure proper localization, we examined PRL-3 membrane targeting using immunofluorescence microscopy with a PRL-3-specific nanobody (57). All functional mutants localized to the plasma membrane in HCT116 cells, consistent with previous reports that used N-terminally tagged constructs (24). A C170S mutant lacking the C-terminal prenylation motif served as a control and exhibited diffuse cytoplasmic localization (**Supplemental Figure 2**), confirming the assay’s sensitivity.

### PRL-3 enhances ALL leukemogenesis and penetrance in a phosphatase-independent manner

To investigate the in vivo relevance of PRL-3’s catalytic and non-catalytic functions, we used a zebrafish model of T-cell Acute Lymphoblastic Leukemia (ALL) driven by *rag2:Myc* overexpression (74). Previous studies have shown that PRL-3 accelerates disease in this model, but it is unclear which of its functions is responsible for these effects (83,84). To confirm compatibility with our zebrafish models, we aligned human and zebrafish PRL-3 and CNNM sequences. PRL-3 is 87% identical between species, and the CNNM CBS domains are 71– 82% identical, with all residues required for binding and mutagenesis fully conserved (**Supplemental Figure 3**). These findings support the use of human PRL-3 mutants to probe functional mechanisms in zebrafish. To generate transgenic animals, embryos were injected with *rag2:Myc* and *rag2:mCherry* constructs to initiate and visualize leukemia, along with either *rag2:mCherry* control or *rag2:PRL-3* variant constructs (WT, C104D, or R138E).

We found both PRL-3 WT and the phosphatase-dead C104D mutant significantly increased leukemia penetrance compared to mCherry controls (18.2% and 21.6% vs. 8.9%; p ≤ 0.044), whereas the CNNM-binding deficient R138E mutant had no effect (*p*=0.997; **Figure 3A–B**). By 20 days post-fertilization (dpf), animals expressing PRL-3 WT or C104D showed significantly higher mCherry signal in the thymus and kidney head than either control and R138E animals (*p*≤0.0038, **Figure 3C-D**), indicating accelerated disease onset. In this model, leukemia originates in the thymus and enters into blood circulation over time (**Supplemental Figure 4**). Circulating ALL cells appeared significantly earlier in PRL-3 WT and C104D (*p*≤0.0057), but not in those expressing R138E (*p*=0.8202, **Figure 3E-F**).

**Figure 3.**
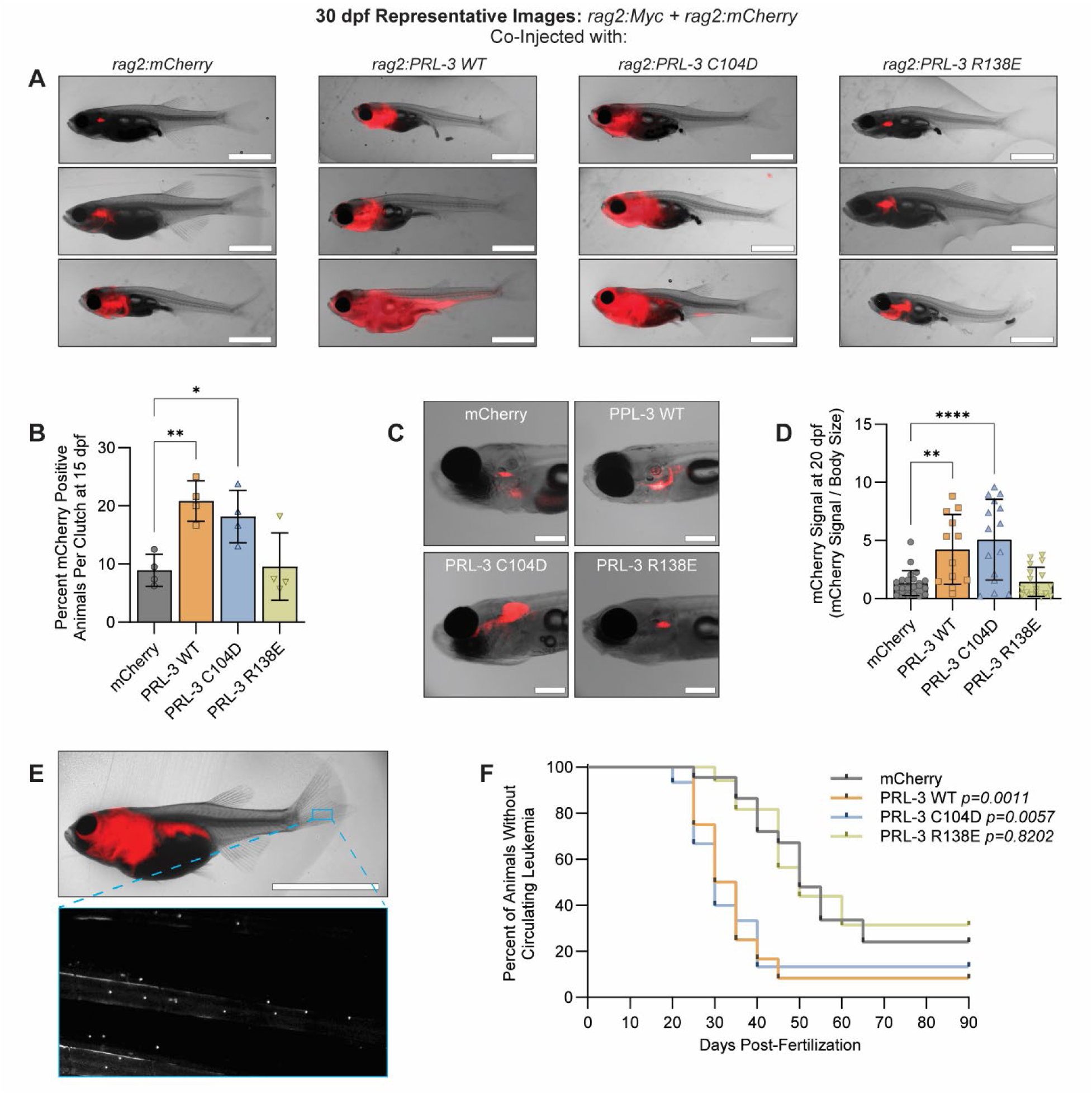
PRL-3 promotes T-cell acute lymphoblastic leukemia (T-ALL)_ progression in vivo through a phosphatase-independent mechanism. **(A)** Representative images of three animals per group at 30 days post-fertilization (dpf) with T-ALL. Scale bars, 2.5 mm. **(B)** Leukemia penetrance quantified as percentage of animals per clutch with detectable mCherry signal at 15 dpf. Each data point represents an individual clutch of at least 25 animals. **(C)** Representative animals at 20 dpf showing mCherry-positive blasts localized to the thymus and kidney head (marrow) regions. Scale bars, 0.5 mm. **(D)** Quantification of mCherry signal relative to body size at 20 dpf. Each data point represents an individual animal; data reflect at least four clutches per group. **(E)** Representative image of a 60 dpf animal with circulating T-ALL. The inset shows mCherry-positive cells within the caudal fin vasculature. Scale bar, 5 mm. **(F)** Kaplan-Meier curve showing time to detectable circulating leukemia, with assessments every 5 days. Data are pooled from at least four clutches per group. Error bars indicate standard deviation. Statistical significance was determined using one-way ANOVA with Tukey correction (B, D) or Kaplan-Meier analysis with log-rank test (F). *, *p*<0.05; **, *p*<0.01; ***, *p*<0.001; ****, *p*<0.0001.

Molecular profiling of isolated leukemia cells confirmed expression of lymphoid (*rag1* and *rag2*) and T-cell specific (*lck* and tcr*β-C2*) markers, with no significant upregulation of B-cell genes (*pax5*, *igD-VH1*, and *igM-VH1*), indicating that PRL-3 did not alter leukemia lineage identity (**Supplemental Figure 5A**). Notably, PRL-3 WT and C104D samples had reduced expression of *tcrβ-C2* compared to the control and R138E groups (*p*≤0.007), suggesting a possible arrest at an earlier developmental state. Expression of *Myc* and *PRL-3* transgenes were comparable across groups (**Supplemental Figure 5B**), confirming that the observed effects are not due to differential transgene expression. Furthermore, we observed no significant differences in blast morphology, proliferation, or viability across conditions **(Supplemental Figures 6-7**), suggesting PRL-3’s impact on leukemia progression is not driven by changes in these parameters.

These results demonstrate that PRL-3 promotes leukemogenesis and accelerates leukemia dissemination in vivo through a phosphatase-independent mechanism. We next asked whether this function similarly contributes to cancer progression in other tumor types, particularly in solid tumors.

### PRL-3 increases RMS tumor size but not tumor initiation, in a phosphatase-independent manner

To asses which function of PRL-3 enhances solid tumor progression, we used a transgenic zebrafish model of rhabdomyosarcoma (RMS) driven by rag2:KRAS^G12D^ (73), which enables real-time visualization of tumor initiation and growth. Embryos were injected with *rag2:KRAS^G12D^* and *rag2:mCherry*, along with either mCherry control or the PRL-3 variant constructs. This model has been used extensively to study RMS biology, and a recent study identified a clinical association between PRL-3 and human RMS progression (85), further supporting its relevance.

PRL-3 expression did not significantly impact tumor initiation, as 7.73-16.68% of animals per clutch developed mCherry-positive nodules by 15 dpf (*p*≥0.656), and zebrafish developed an average of 1.2 – 1.7 tumors per animal across all groups (*p*=0.809) (**Figure 4A-C, Supplemental Figure 8A-C**). As a positive control for enhanced tumorigenesis, the *p53*^-/-^ strain (76) had significantly higher tumor incidence (average 27.58% per clutch, *p*=0.0117) as well as more tumors per animal (average 2.79 tumors per animal, *p*<0.0001, **Supplemental Figure 9**), confirming the system’s sensitivity. These findings suggest that PRL-3 does not affect tumor initiation in this RMS model.

**Figure 4.**
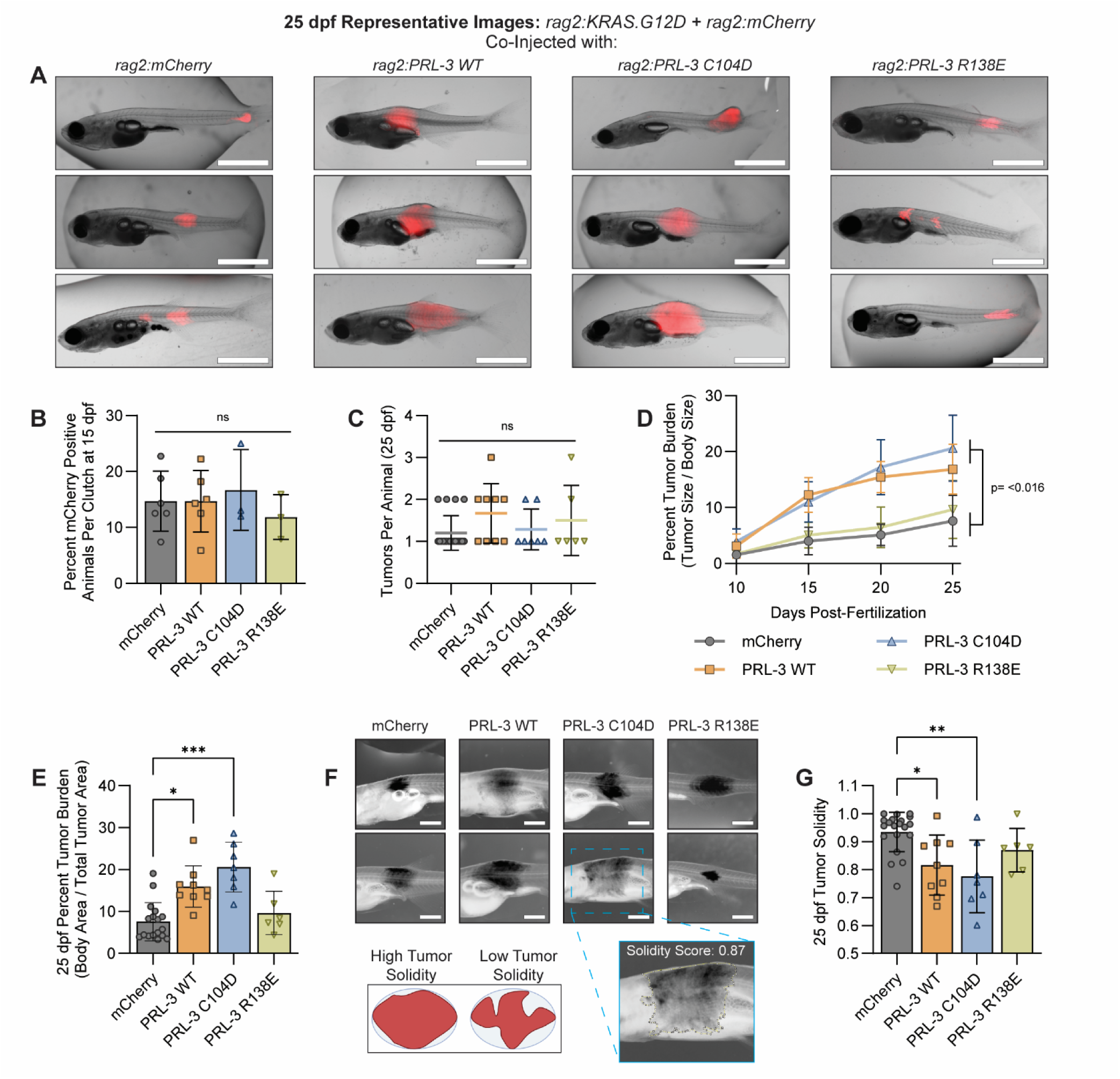
PRL-3 promotes rhabdomyosarcoma progression in vivo through a phosphatase-independent mechanism. **(A)** Representative images of three animals per group at 25 days post-fertilization (dpf), co-injected with *rag2:KRAS^G12D^*, *rag2:mCherry*, and either *rag2:mCherry* control or *rag2:PRL-3* variant constructs, as indicated. Scale bars, 2.5 mm. **(B)** Quantification of tumor penetrance at 15 dpf, shown as a percentage of larvae per clutch with detectable mCherry signal. Each data point represents one clutch of at least 25 animals. **(C)** Number of tumors per animal at 25 dpf. Each data point represents an individual animal. Zebrafish with tumor burden greater than 50% of their total body area were excluded from quantification. **(D)** Longitudinal quantification of tumor burden, calculated as tumor area relative to body size. Only the largest tumor was selected for quantification from larvae with multiple tumors. **(E)** Quantification of tumor burden at the 25 dpf time point. Each point represents an individual zebrafish; data are pooled from ≥3 clutches per group. **(F)** Representative images of two animals per group used for solidity analysis. Image contrast is inverted for clarity. A schematic shows scoring criteria for high vs low tumor solidity. Scale bars, 1 mm. **(G)** Tumor solidity at 25 dpf. Each point represents an individual tumor; only tumors along the body axis were included in the analysis to avoid variability from tumors with divergent morphology (e.g., dorsal fin or jaw). For all, error bars represent standard deviation. Statistical significance was assessed using one-way ANOVA with Tukey correction (B, C, E, G) or two-way ANOVA with Tukey correction (D). *, *p*<0.05; **, *p*<0.01; ***, *p*<0.001.

Despite not altering tumorigenesis, PRL-3 WT and the phosphatase-dead C104D mutant increased tumor size, quantified as percent tumor burden (tumor size/body size), with both averaging ≥15.98% compared to 7.59% in the mCherry controls (*p*≤0.025). The CNNM-binding deficient R138E mutant did not alter tumor size (*p*>0.991, **Figure 4D-E**). The A111S “enhanced-phosphatase” mutant further increased tumor burden (average 30.54%, *p*<0.0001), whereas A111S+R138E and C104E mutants showed no significant effects (≤11.50%, *p*≥0.224) (**Supplemental Figure 8D-E).** These findings suggest that while elevated phosphatase activity may enhance PRL-3’s effects on tumor growth, CNNM binding remains essential for this phenotype. Additionally, tumor solidity, a morphological feature associated with invasiveness (86–88), was significantly reduced in PRL-3 WT and C104D tumors compared to control (p≤0.013), suggesting a more diffuse growth pattern. This reduction in tumor solidity was recapitulated by A111S (*p*=0.002) but not R138E, C104E, or A111S+R138E mutants (*p*≥0.419, **Figure 4F-G, Supplemental Figure 8F**).

All RMS tumors, regardless of group, showed significantly elevated expression of *cdh15* and *myog* compared to normal muscle (*p*<0.0001), consistent with a poorly differentiated state (73). Only the C104D group exhibited increased *desma* expression (*p*=0.009), which may indicate a differentiation arrest (89–91). Other muscle lineage markers (*pax7b*, *mylf5*) were unchanged. Expression levels of *KRAS^G12D^* and *PTP4A3* were consistent across the experimental groups (**Supplemental Figure 10**), ruling out differential transgene expression as a confounding factor. Representative H&E sections from RMS tumors are provided in **Supplemental Figure 11**.

Our results indicate that PRL-3 promotes RMS tumor growth and alters morphology through a phosphatase-independent mechanism, likely driven by CNNM binding. To further define the cellular basis of PRL-3 mediated tumor progression, we next evaluated its role in self-renewal, anchorage-independent growth, migration, and invasion in vitro. These phenotypes have been previously linked to PRL-3 expression in various cancer types (16,18,92,93), though their molecular basis remains unclear.

### PRL-3 enhances self-renewal and anchorage-independent growth in a phosphatase-independent manner

To broadly investigate the role of PRL-3 in cancer aggressiveness, we generated doxycycline-inducible cell lines in three cancer types: Jurkat (T-cell acute lymphoblastic leukemia), RD (embryonal rhabdomyosarcoma), and HCT116 (colon carcinoma). Western blot analysis confirmed doxycycline-dependent expression of PRL-3 and its mutants in all three lines (**Figure 5A-C**).

**Figure 5.**
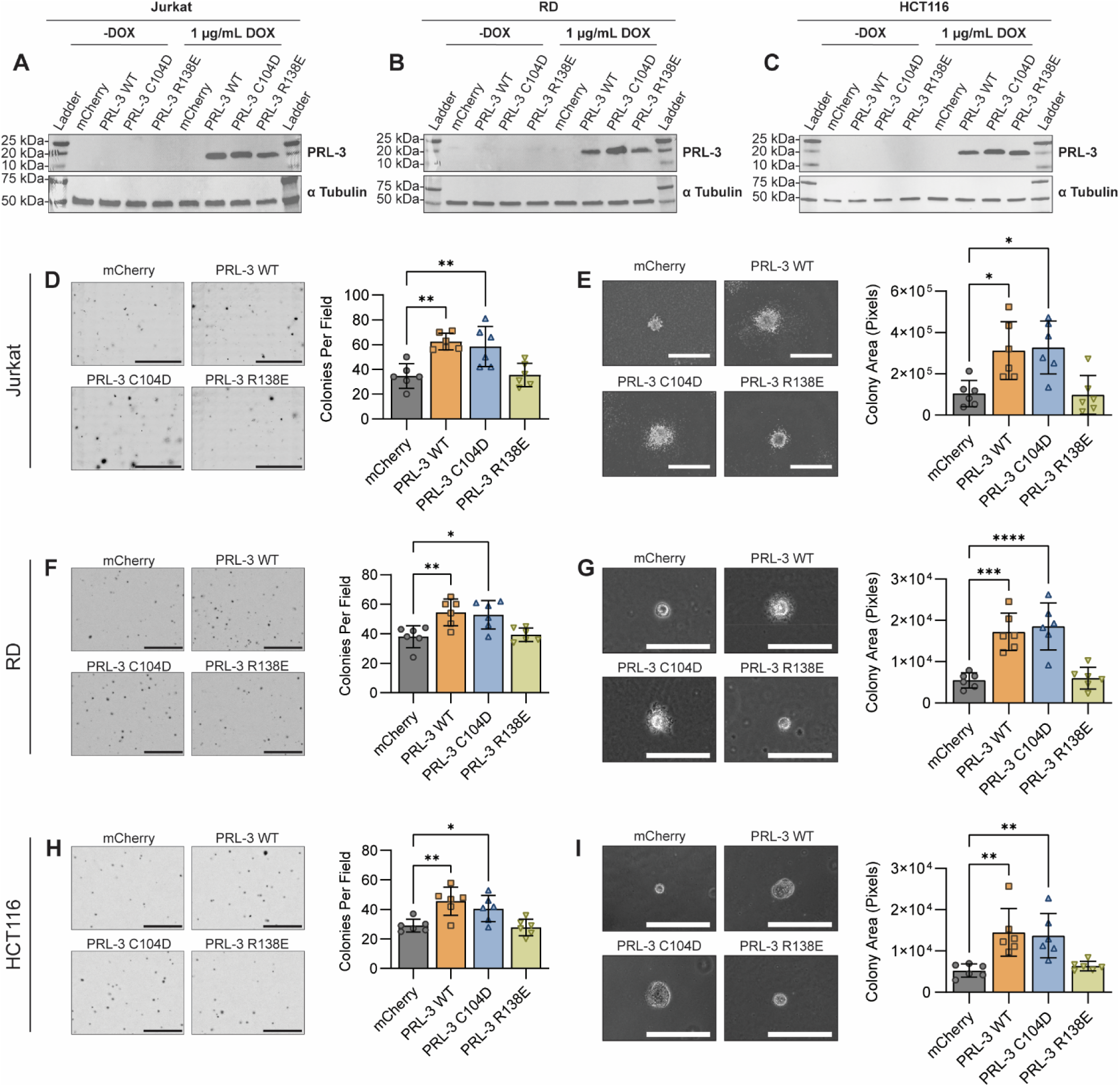
PRL-3 enhances colony formation and anchorage-independent growth in a phosphatase-independent manner. (A-C) Western blots showing the inducible overexpression of mCherry (control), PRL-3 wild-type (WT), and PRL-3 mutants in Jurkat (**A**), RD (**B**), and HCT116 (**C**) cell lines following doxycycline treatment.α-Tubulin serves as a loading control. Blots are representative of at least two independent experiments. **(D)** Representative images and quantification of Jurkat colonies cultured in MethoCult medium. Colonies >50 µm in diameter were counted per field; data points represent the average of two technical replicates, pooled from three independent experiments. Scale bar, 10 mm. **(E)** Representative images and quantification of Jurkat colonies size for each condition. Colony area (pixels) was quantified from 25 randomly selected colonies per well. Data are averaged per condition across three experiments. Scale bar, 1000 µm. **(F, H)** Representative images and quantification of RD **(F)** and HCT116 **(H)** colonies grown in soft agar. Colonies >50 µm in diameter were counted per field. Data points represent the average results of two technical replicates, pooled from three independent experiments. **(G, I)** Representative images and quantification of individual RD **(G)** and HCT116 **(I)** colony sizes. Colony areas were quantified from 10 randomly selected colonies per well. Data are shown as average size per condition across three independent experiments. Scale bars, 100 µm (G) and 400 µm (I). For all, error bars represent standard deviation. Statistical significance was assessed using one-way ANOVA with Tukey correction. *, *p*<0.05; **, *p*<0.01; ***, *p*<0.001; ****, *p*<0.0001

To test the impacts of PRL-3 on the self-renewal capacity of suspension cells, we performed a MethoCult-based colony formation assay (CFA) using Jurkat cells. This assay evaluates the ability of cells to proliferate clonally, serving as a proxy for increased disease aggressiveness and potential for relapse (94,95). Overexpression of PRL-3 WT and the phosphatase-dead C104D mutant significantly increased colony formation of (≥58.5 colonies per field) compared to both the mCherry control and the CNNM-binding deficient R138E mutant (≤35.5 colonies per field; *p*≤0.007, **Figure 5D**). Colonies in the PRL-3 WT and C104D groups were also significantly larger (*p*≤0.019), with a more diffuse morphology, suggestive of increased outward cell migration from the colony core (**Figure 5E**).

The soft agar colony formation assay is a widely used in vitro method to evaluate anchorage-independent growth of adherent cell lines as an indicator of enhanced aggressiveness (96,97). In RD cells, PRL-3 WT and C104D significantly increased colony numbers (≥52.8 colonies per field) and colony size (*p*≤0.0005) relative to the mCherry control and the R138E mutant (≤39.33 colonies per field; *p*≤0.01; **Figure 5F-G**). Similarly, HCT116 cells expressing PRL-3 WT and C104D formed more colonies (≥41.83 colonies per field) than control and R138E groups (≤28.17 colonies per field, *p*≤0.038) and exhibited significantly larger colony sizes (*p*≤0.024; **Figure 5H-I**).

While larger colonies may suggest enhanced cell proliferation, growth rate analysis revealed no significant differences between the experimental groups (**Supplemental Figure 12**), indicating that PRL-3-driven effects on colony formation are not due to increased cell division. Interestingly, across all three cells lines, PRL-3 WT and C104D consistently produced colonies with a more diffuse morphology, suggesting increased motility. Building on these observations, we next performed transwell assays to directly assess the impact of PRL-3 on cell migration and invasion.

### PRL-3 enhances cell migration and invasion in vitro in a phosphatase-independent manner

We used transwell assays to assess which function of PRL-3 promotes cell motility and invasion. Jurkat cells, which have limited invasive capacity, were analyzed in uncoated transwell migration assays, while RD and HCT116 were assessed using Matrigel-coated invasion chambers to evaluate matrix penetration. These assays measure the ability of cells to respond to chemotactic cues, reorganize the cytoskeleton, and, in the case of invasion, activate matrix-degrading enzymes (98,99).

In Jurkat cells, expression of PRL-3 WT and the phosphatase-dead C104D mutant significantly increased migration (≥1.28-fold compared to the mCherry control; *p*≤0.013), whereas the CNNM-binding deficient R138E mutant had no effect (*p*=0.9175) (**Figure 6A**). In RD and HCT116 invasion assays, PRL-3 WT and C104D similary increased invasion (≥1.627-fold in RD, ≥1.44-fold in HCT116) compared to mCherry (*p*≤0.014 and *p*≤0.039, respectively), while R138E failed to enhance invasion (RD: *p*=0.973; HCT116: *p*=0.945; **Figure 6B-C**). These results demonstrate that PRL-3 promotes migration and invasion through a mechanism independent of its phosphatase activity. Although we cannot exclude the possibility that the R138E mutation disrupts interactions with proteins other than CNNMs, its consistent loss of function across models suggests that this mutated interface is critical for PRL-3’s oncogenic effects. Given prior studies implicating the R138 region in CNNM binding, these results are consistent with a model in which the PRL-3:CNNM interaction is a key driver of aggressive phenotypes. To further explore this mechanism, we next evaluated whether existing PRL-3 inhibitors can disrupt the PRL-3:CNNM interaction, and we developed a FRET-based assay that can serve as a platform for future screening of compounds targeting this interface.

**Figure 6.**
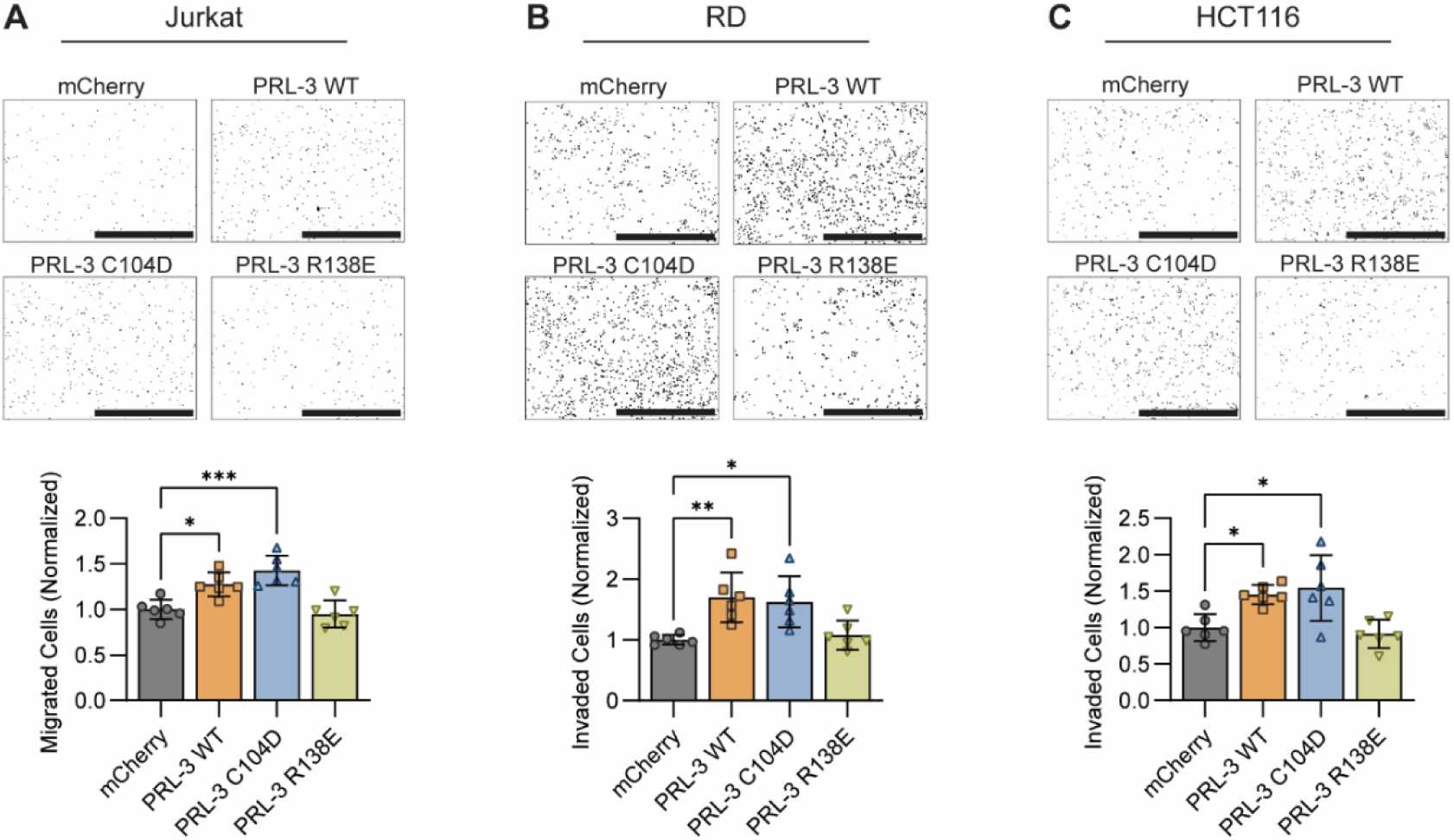
PRL-3 enhances migration and invasion through a phosphatase-independent mechanism. **(A)** Representative images and quantification of Jurkat cells that migrated through uncoated transwells and adhered to poly-D-lysine-coated plates, stained with Hoechst. Scale bar, 1mm. **(B, C)** Representative images and quantification of RD **(B)** and HCT116 **(C)** cells that invaded through Matrigel-coated transwells. Scale bars, 1 mm. For all, quantification represents migrated or invaded cell counts normalized to the mCherry control group. Each data point represents an individual transwell; data are pooled from three independent experiments. Error bars indicate standard deviation. Statistical significance was determined using one-way ANOVA with Tukey correction. *, *p*<0.05; **, *p*<0.01; ***, *p*<0.001; ****, *p*<0.0001.

### Development of a High-Throughput Screen for PRL:CNNM Interaction Inhibitors

To enable high-throughput screening for inhibitors of the PRL-3:CNNM interaction, we developed an in vitro fluorescence resonance energy transfer (FRET)-based assay. FRET occurs when the donor and acceptor fluorophores are brought into close proximity, here, through binding of PRL-3 and the CBS domain of CNNM, resulting in acceptor emission upon donor excitation. Disruption of this interaction reduces the FRET signal.

We engineered recombinant proteins by fusing large Stokes shift mGFP (LSSmGFP) to the N-terminus of full-length PRL-3 (FRET donor) and mScarlet3 to the C-terminus of the HA-tagged CBS domain of CNNM3 (FRET acceptor). The purity and fluorescent properties of the proteins are shown in **Supplemental Figure 13**. The C104D mutant was used to prevent phosphatase activity and eliminate the need for reducing agents, ensuring the assay reflected binding rather than catalytic function (31). Both fluorescent fusion proteins retained the ability to co-immunoprecipitate with their untagged binding partners, confirming that the fluorescent tags did not disrupt interaction (**Supplemental Figure 14A**). FRET signal was measured in this assay by detecting HA-CBS-mScarlet3 emission (600 nm) upon excitation of LSSmGFP-PRL-3 at 400 nm (**Figure 7A**) and showed a dose-dependent saturation with increasing concentrations of HA-CBS-mScarlet3 (**Supplemental Figure 14B**) Signal specificity was further confirmed by competition with unlabled HA-CBS, which reduced FRET response, whereas bovine serum albumin (BSA), included to control for non-specific effects such as protein crowding, had no effect (**Supplemental Figure 14C**). These data validate the FRET assay as a specific and functional platform for quantifying PRL-3:CNNM interactions in vitro.

**Figure 7.**
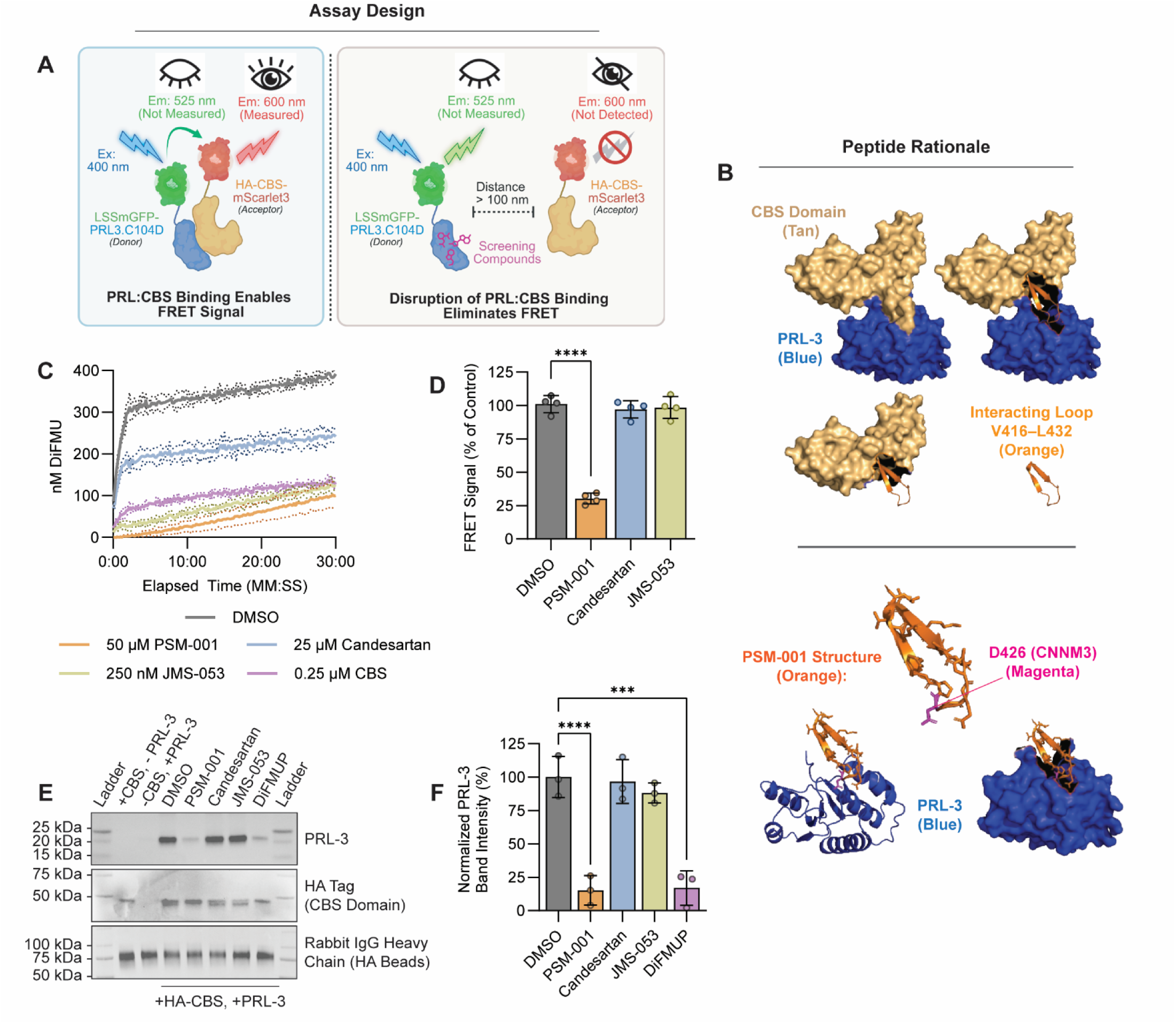
Development of a high-throughput assay to screen for inhibitors of the PRL:CNNM interaction. **(A)** Schematic illustration of the FRET assay design. When LSSmGFP-PRL3.C104D binds HA-CBS-mScarlet3, excitation at 400nm induces emission at 600 nm. Disruption of this interaction increases donor-acceptor distance and abolishes the FRET signal. **(B)** Structural rationale for cyclic peptide design. The PRL-interacting loop of CNNM3 (V416-L432) was cyclized to generate PRL substrate mimics (PSM), with PSM-001 used as a positive control for binding disruption. **(C)** DiFMUP-based phosphatase assay showing PRL-3 activity in the presence of the indicated inhibitors. The purified CBS domain served as an additional control. Data are representative of two independent experiments; dotted lines indicate standard deviation. **(D)** Application of the FRET assay to test PSM-001 and existing PRL-3 inhibitors (candesartan, JMS-053), each at 50 µM. Fluorescence intensity is normalized to DMSO control. Data are pooled from four independent experiments. **(E)** In vitro co-immunoprecipitation of purified PRL-3 and HA-CBS following treatment with 50 µM of the indicated compounds. The blot shown is representative of at three independent experiments. **(F)** Quantification of PRL-3 band intensity in (E), normalized to DMSO control. Data are pooled results from three experiments. Error bars indicate standard deviation; statistical significance was determined by one-way ANOVA and Tukey correction. ****, *p*<0.0001.

As a positive control, we designed cyclic peptides based on the CNNM3 binding interface (residues V416-L432; **Figure 7B and Supplemental Table 3**). Circular dichroism (CD) analysis revealed that one peptide, PSM-001 displayed the most apparent secondary structure, and was the most effective at inhibiting PRL-3 phosphatase activity in vitro (**Supplemental Figure 15**), consistent with its ability to mimic CNNM binding. Based on these properties, PSM-001 was selected as a tool compound for further investigation.

In phosphatase activity assays, PSM-001 and previously characterized PRL-3 inhibitors, candesartan and JMS-053 (KVX-053) (48,51,100,101) each showed varying degrees of catalytic inhibition (**Figure 7C**). However, only PSM-001 significantly reduced the FRET signal (by 40%; *p*<0.0001), while candesartan and JMS-053 had no effect (*p*≥0.823) (**Figure 7D**). These results were corroborated by co-immunoprecipitation assays using non-fluorescent proteins, in which only PSM-001 disrupted PRL-3:CNNM binding (**Figure 7E-F**). These findings demonstrate that PSM-001 can inhibit the PRL-3:CNNM interaction in vitro, while existing PRL-3 inhibitors, despite reducing phosphatase activity, do not disrupt this critical protein-protein interaction.

## Discussion

The oncogenic mechanism of PRL-3 has remained poorly defined despite decades of study. A major barrier to mechanistic insight has been the field’s reliance on the PRL-3 C104S as a loss-of-function tool. Although this mutation was intended to eliminate phosphatase activity, it also alters PRL-3 catalytic site in a way that prevents CNNM binding (34). As a result, prior investigations have been unable to determine whether PRL-3 enhances malignancy through catalytic activity, protein–protein interactions, or both. Our study addresses this limitation using structure-guided PRL-3 mutants, previously validated in vitro, to functionally separate phosphatase activity from CNNM binding (24). Across two in vivo zebrafish cancer models, T-cell acute lymphoblastic leukemia and rhabdomyosarcoma, as well as multiple human cancer cell lines, the phosphatase-dead C104D mutant consistently recapitulated the phenotypes induced by wild-type PRL-3. In contrast, a catalytically active mutant that cannot bind CNNM proteins (R138E) failed to enhance oncogenesis, tumor growth, or dissemination in vivo, and did not promote aggressive phenotypes such as self-renewal, migration, and invasion in vitro. These findings argue that PRL-3 promotes cancer progression through non-catalytic mechanisms, and that its phosphatase activity is dispensable in this context.

These findings have significant implications for the PRL field, which has historically focused on the phosphatase activity of this protein. Considerable effort has gone into identifying substrates and developing phosphatase inhibitors, yet our data suggest that these strategies may not effectively block PRL-3’s cancer-relevant functions. Instead, protein–protein interactions, particularly with CNNM magnesium transporters, appear to be the critical mediators of its oncogenic activity. This shifts focus away from substrate dephosphorylation as the primary mechanism and suggests that therapeutic efforts should target PRL-3’s binding interfaces rather than its catalytic domain. To support future work, we developed a FRET-based in vitro assay using purified fluorescent proteins to quantify PRL:CNNM binding and evaluate potential inhibitors. We validated a cyclic peptide as a tool compound capable of disrupting this interaction. While not intended as a therapeutic lead, this peptide provides a positive control for future structure-guided inhibitor development.

The biological consequences of PRL-3:CNNM binding are still being defined, but several studies suggest this interaction may regulate magnesium homeostasis in ways that contribute to cancer progression. PRL binding inhibits CNNM-mediated magnesium export and enhances TRPM7-driven import, altering intracellular magnesium levels, although the direct contribution of these changes to tumor progression remains unclear (42). Our findings support CNNM binding as a functionally important activity of PRL-3, which occurs through its highly conserved catalytic site (102), making phosphatase activity and binding mutually exclusive. This structural constraint may explain why PRL-3 has retained a catalytically competent active site rather than evolving into a true pseudophosphatase. We speculate that, in the context of cancer, the phosphatase domain may function primarily to regulate protein-protein interactions rather than substrate dephosphorylation. In support of this model, we found that the A111S mutant, previously shown to increase catalytic turnover (27,82), further enhanced tumor size in the RMS model. However, this effect was abolished when combined with the R138E mutation, suggesting that CNNM binding is still required for this phenotype. One possible explanation is that increased turnover accelerates PRL-3’s release from the phosphocysteine intermediate state, allowing more frequent CNNM binding events (34). While speculative, this finding raises the possibility that PRL-3’s catalytic activity fine-tunes, rather than drives, its oncogenic function through the regulation of protein-protein interactions.

While our data strongly supports CNNM binding as the primary mechanism underlying PRL-3’s oncogenic effects, we interpret the R138E phenotype with appropriate caution. Although this mutation disrupts CNNM interaction in vitro, it may also interfere with other, uncharacterized protein partners. Further studies are needed to identify the full repertoire of PRL-3 binding interactions and to determine whether CNNMs act as obligate effectors or part of a broader binding network.

This study also raises important questions about other PRL family members, such as PRL-1 and PRL-2, which share high structural similarity and may function through analogous mechanisms. Moreover, magnesium homeostasis is increasingly recognized as a regulator of cancer cell behavior (103), yet the downstream consequences of altered magnesium flux remain poorly understood. Our findings highlight the need to revisit longstanding assumptions about the functional roles of PRL proteins and to consider non-catalytic mechanisms when evaluating their contributions to cancer and other biological processes.

In summary, our study demonstrates that PRL-3 promotes cancer progression through a mechanism independent of its phosphatase activity and instead relies on protein-protein interactions, likely with CNNM transporters. These results challenge substrate-centric models of PRL-3 function and support a shift toward therapeutic strategies that target its binding interfaces rather than its catalytic activity.

## Acknowledgments

This work was supported by funding from the National Cancer Institute (R37CA227656) and the Kentucky Pediatric Cancer Research Trust Fund (both to J.S.B.). J.T.J. was supported by a University of Kentucky Lyman T. Johnson Fellowship. This work was further supported by the Markey Cancer Center’s Flow Cytometry & Immune Monitoring and Biospecimen Procurement and Translational Pathology Shared Resources (P30CA177558). Portions of the editing process were supported using Grammarly AI for language refinement. We would like to thank Dr. Yelena Chernyavskaya for her support with the molecular biology techniques and zebrafish experiments, as well as Eva Aldarondo, Chase Yost, and Lexi Baker for their support with zebrafish husbandry and maintenance. Furthermore, we would like to thank Ty Cheatham for his assistance with protein purification and assay optimization.

---

**Supplemental Figure 1.**
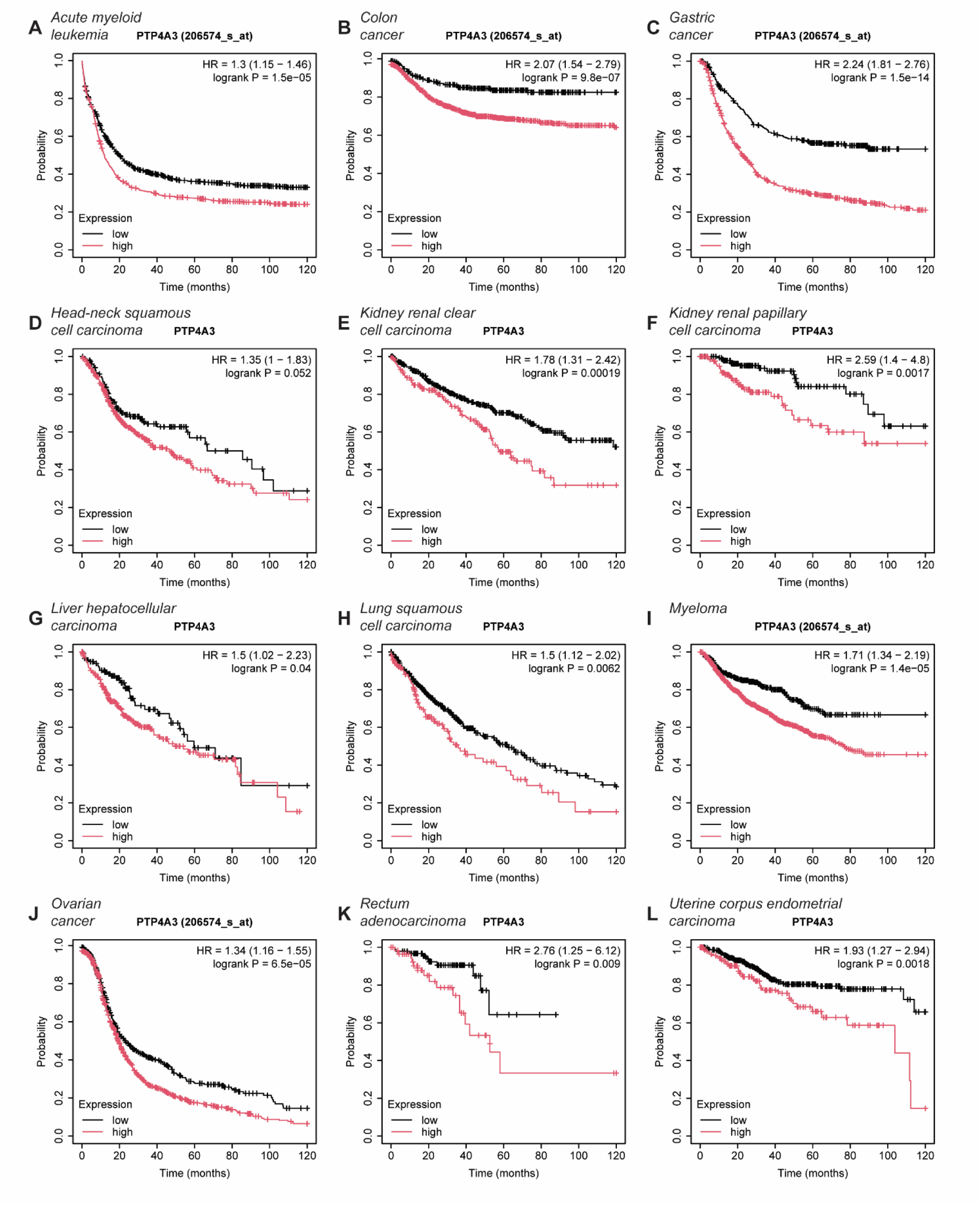
High PRL-3/PTP4A3 expression correlates with a poor patient prognosis. Kaplan-Meier analysis of overall survival in **(A)** acute myeloid leukemia**, (B)** colon cancer, **(C)** gastric cancer, **(D)** head-neck squamous cell carcinoma, **(E)** kidney renal clear cell carcinoma, **(F)** kidney renal papillary cell carcinoma, **(G)** liver hepatocellular carcinoma, **(H)** lung squamous cell carcinoma, **(I)** myeloma, **(J)** ovarian cancer, **(K)** rectum adenocarcinoma, and **(L)** uterine corpus endometrial carcinoma. Sample cohorts are split into PTP4A3 low (black) and high (red) by the median. Hazard ratios were calculated for the PTP4A3 high groups. Data shown is derived from the online tool KM-plotter and pools from microarray (PTP4A3 [206574_s_at]) and RNAseq (PTP4A3) data sets.

**Supplemental Figure 2.**
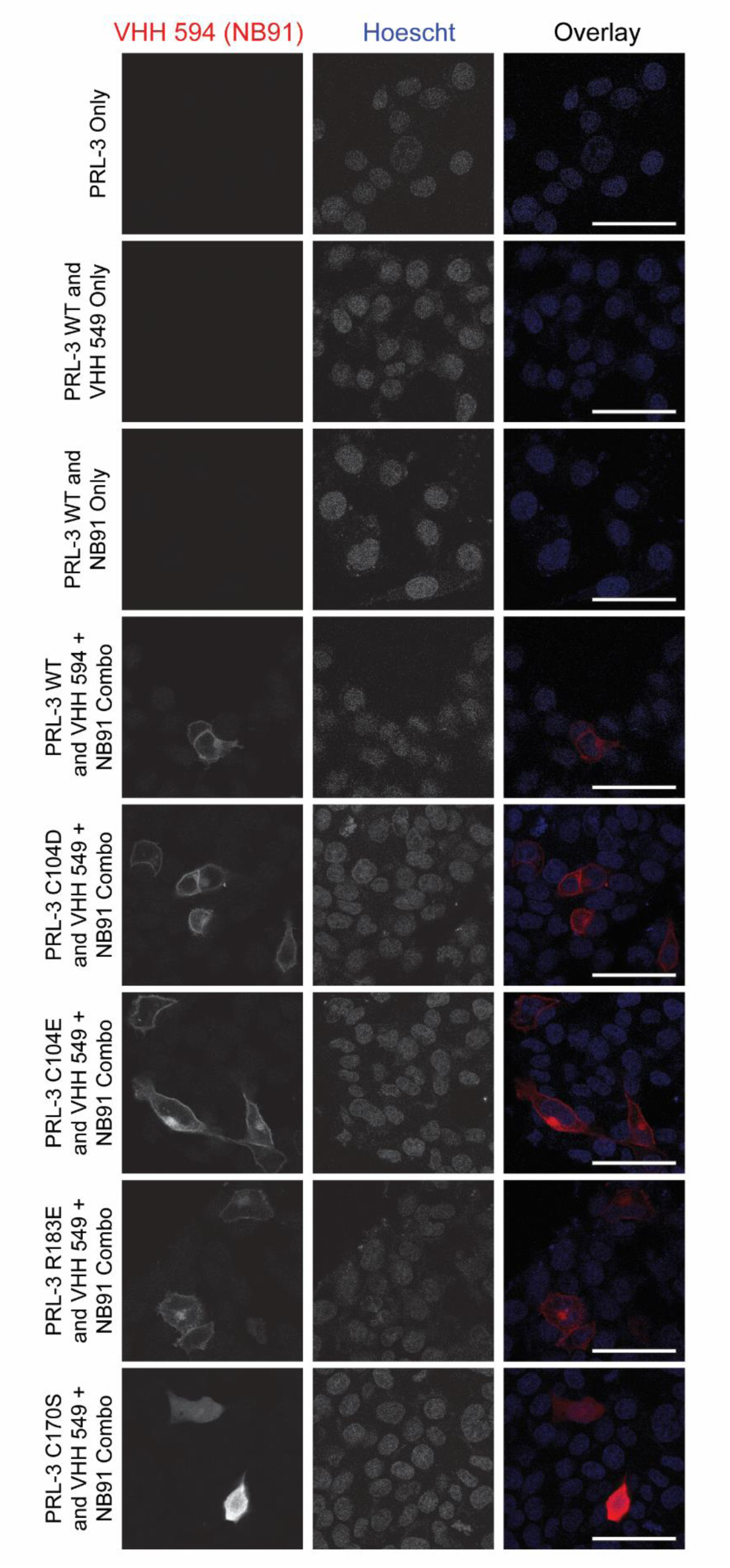
PRL-3 point mutations retain the localization patterns of the wild-type protein. Immunofluorescence (IF) images of HCT116 cells transfected with PRL-3 WT or mutants to assess subcellular localization. Nuclei were visualized with Hoechst 33342 (blue). PRL-3 was detected using a PRL-3-specific nanobody (NB91) and visualized with an anti-alpaca AlexaFluor 594 (VHH 594, red). Scale bar, 20 µm.

**Supplemental Figure 3.**
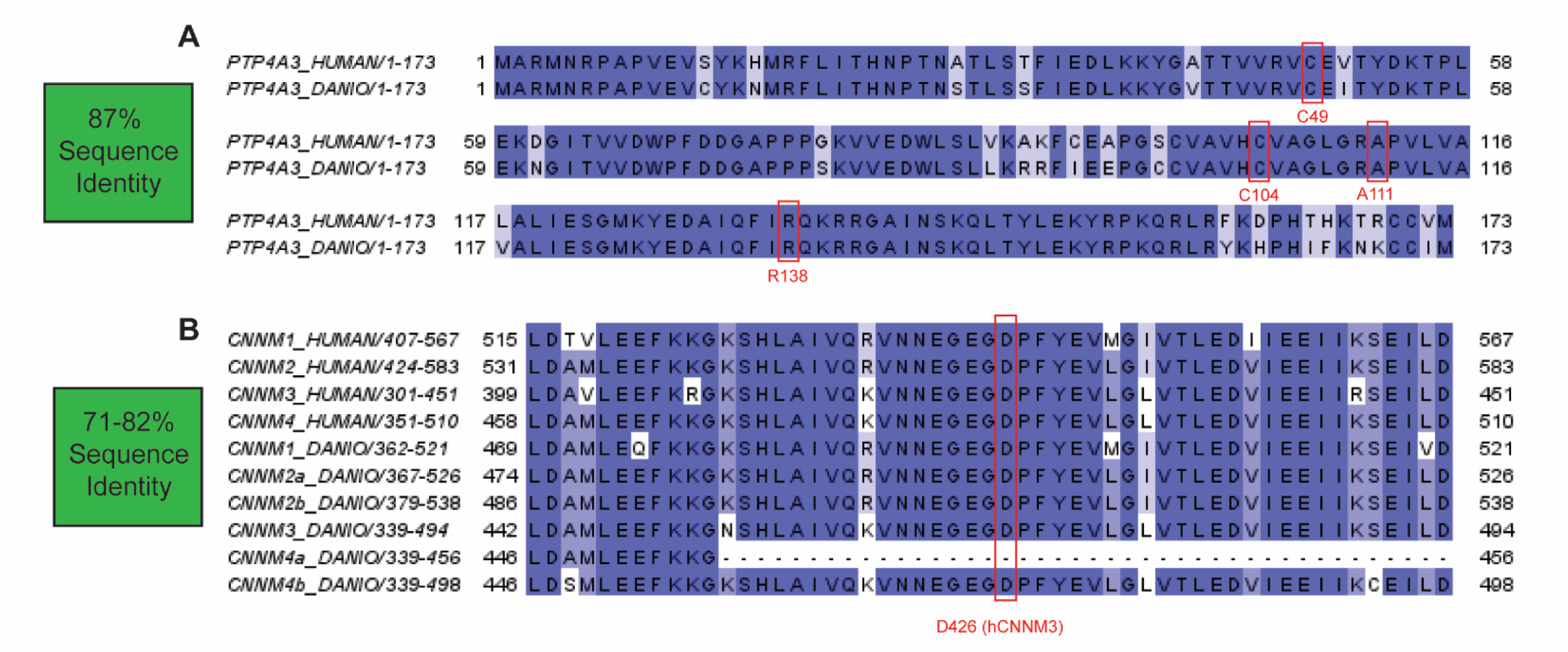
Sequence alignments show that PRL-3 and the CBS domain of CNNMs are highly conserved across humans and zebrafish. **(A)** Protein sequence alignment of human and zebrafish (*Danio rerio*) PRL-3 (PTP4A3) with residues of interest in red, including the C104 catalytic site and R138, which is critical for CBS-domain binding. **(B)** Protein sequence alignment of the CBS domains of human and zebrafish CNNM proteins. Mutation of D426, shown in red, has been shown to reduce PRL-3 binding.

**Supplemental Figure 4.**
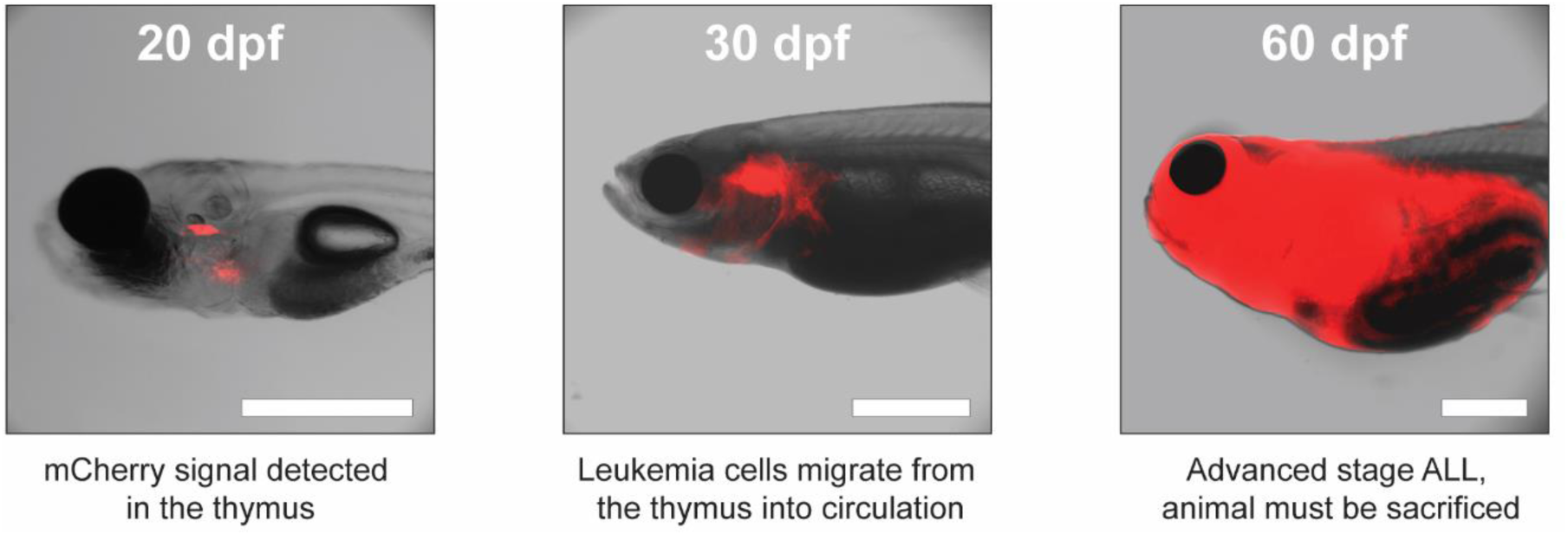
Leukemia in the zebrafish T-ALL model initiates in the thymus and disseminates systemically. Representative images of a single rag2:myc;rag2:mCherry transgenic zebrafish were captured at 20, 30, and 60 days post-fertilization (dpf) to monitor leukemia progression over time. mCherry-positive leukemia cells first accumulate in the thymus and later spread into the circulation. Scale bar, ∼1 mm.

**Supplemental Figure 5.**
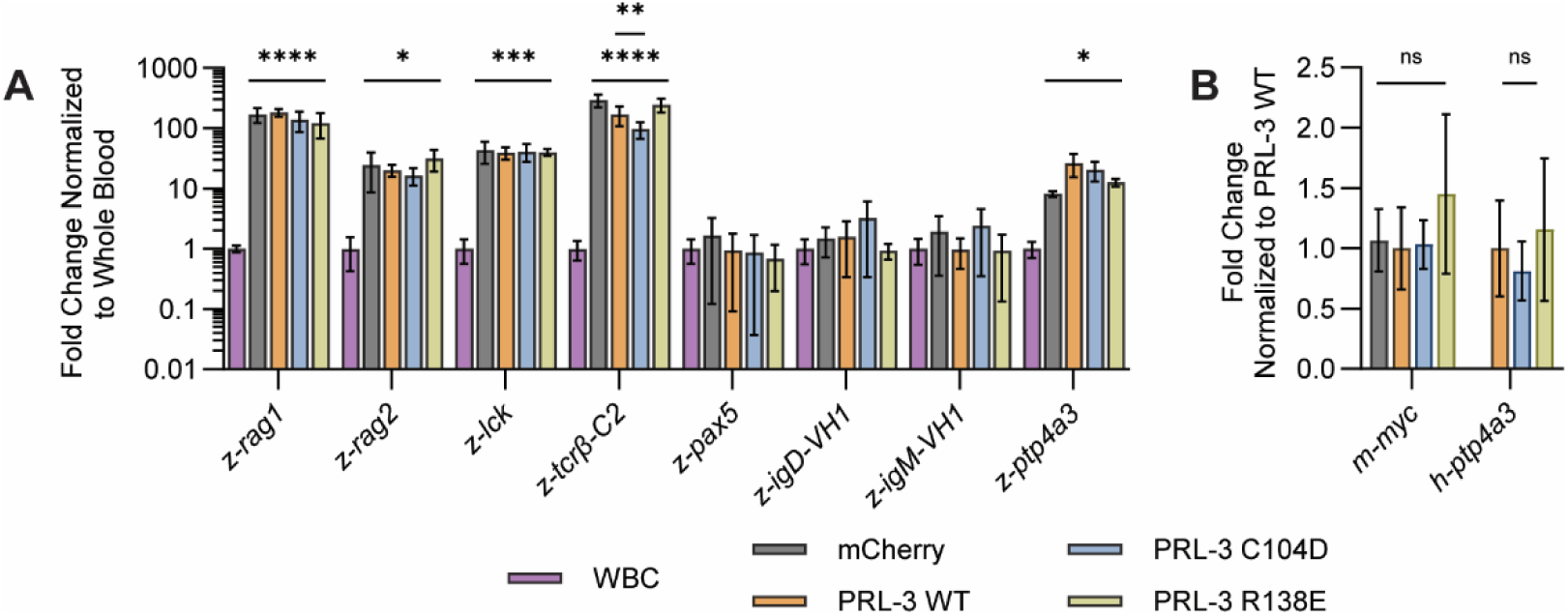
The overexpression of PRL-3 mutants has a minimal effect on ALL gene signatures. **(A)** RT-qPCR analysis of endogenous genes involved in lymphoid, T cell, or B cell specification in the indicated ALL samples, normalized to expression in non-cancerous whole blood (WBC) **(B)** RT-qPCR analysis results for injected transgenes normalized to the PRL-3 WT group. Each analysis was performed with samples extracted from six animals per condition. Error bars represent standard deviation. Statistical significance was calculated using GraphPad Prism with an ordinary one-way ANOVA and Tukey’s multiple comparisons test. *, *p*<0.05; **, *p*<0.01; ***, *p*<0.001; ****, *p*<0.0001.

**Supplemental Figure 6.**
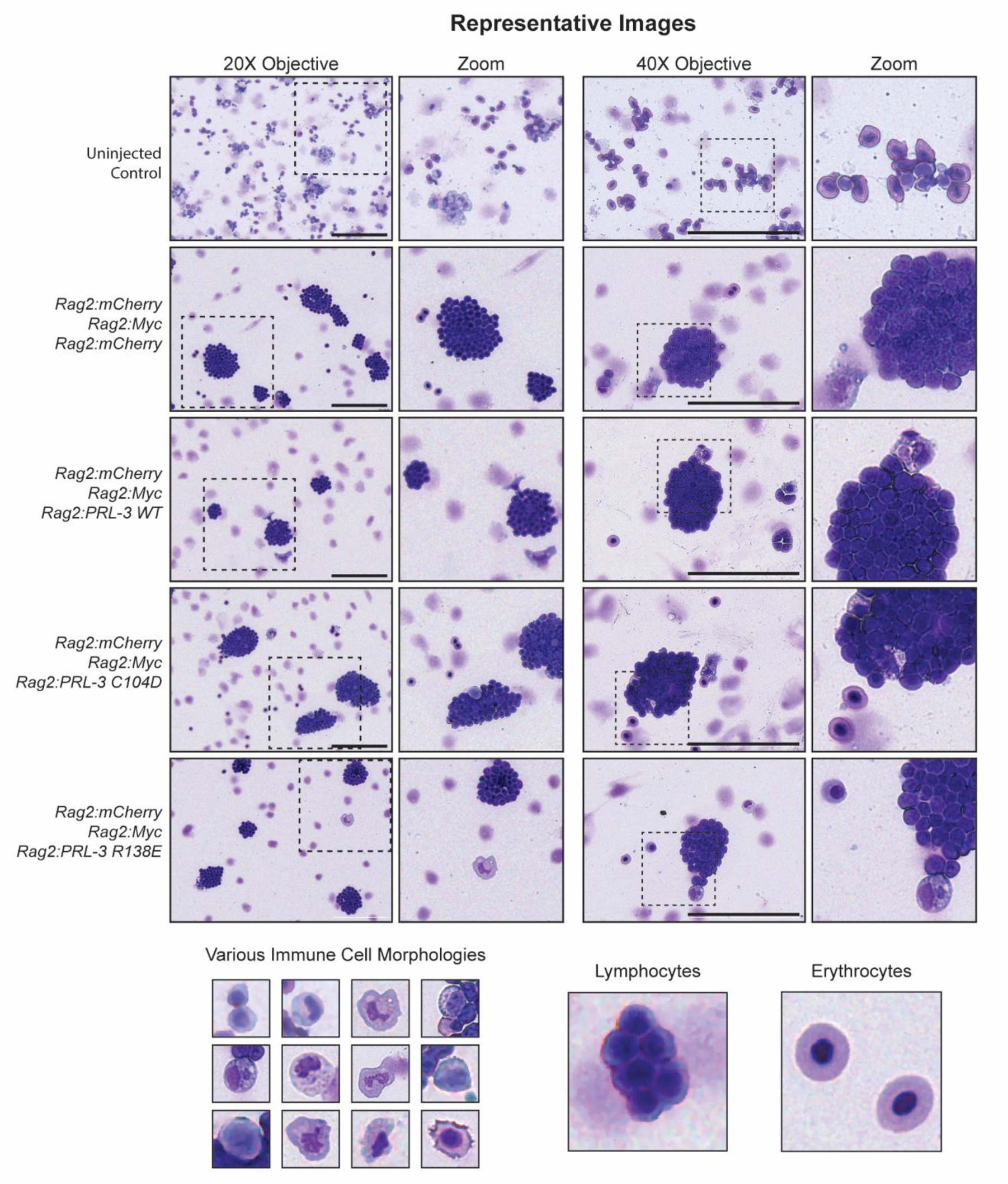
Representative May-Grünwald-Giemsa staining of zebrafish ALL samples. Zebrafish acute lymphoblastic leukemia (ALL) samples from each experimental group were stained with May-Grünwald-Giemsa to assess cellular morphology. Images shown are representative of at least two animals per group. Scale bar, 100 µm.

**Supplemental Figure 7.**
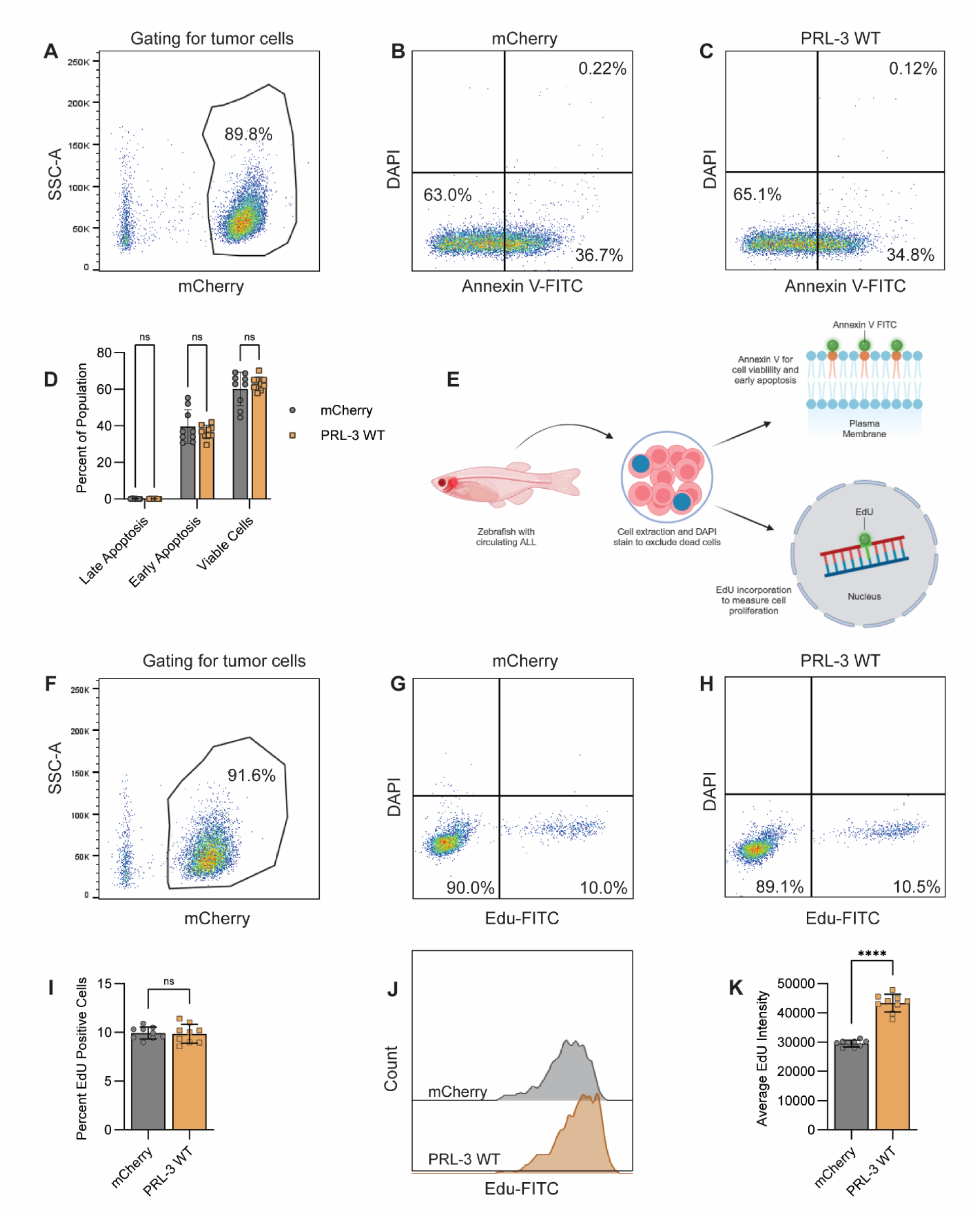
Annexin V and EdU analysis of ALL samples. **(A)** Representative plot for gating of mCherry-positive tumor cells for Annexin V staining. **(B, C)** Representative Annexin V staining plots for mCherry control and PRL-3 WT groups **(D)** Quantification of Annexin V-positive cells, data points represent three animals per condition. **(E)** Schematic illustrating the methodology for Annexin V and EdU assays. **(F)** Representative plot for gating of mCherry-positive tumor cells for EdU analysis. **(G, H)** Representative EdU incorporation plots for mCherry control and PRL-3 WT groups. **(I)** Quantification of EdU-positive cells. Data points are the pooled reads from three animals per group. **(J)** Representative EdU signal intensity plots for mCherry and PRL-3 WT groups. **(K)** Quantification of the EdU signal intensity per condition. Data points are the pooled reads from three animals per group. Error bars represent standard deviation; significance was determined using an unpaired *t*-test. ****, *p*<0.0001.

**Supplemental Figure 8.**
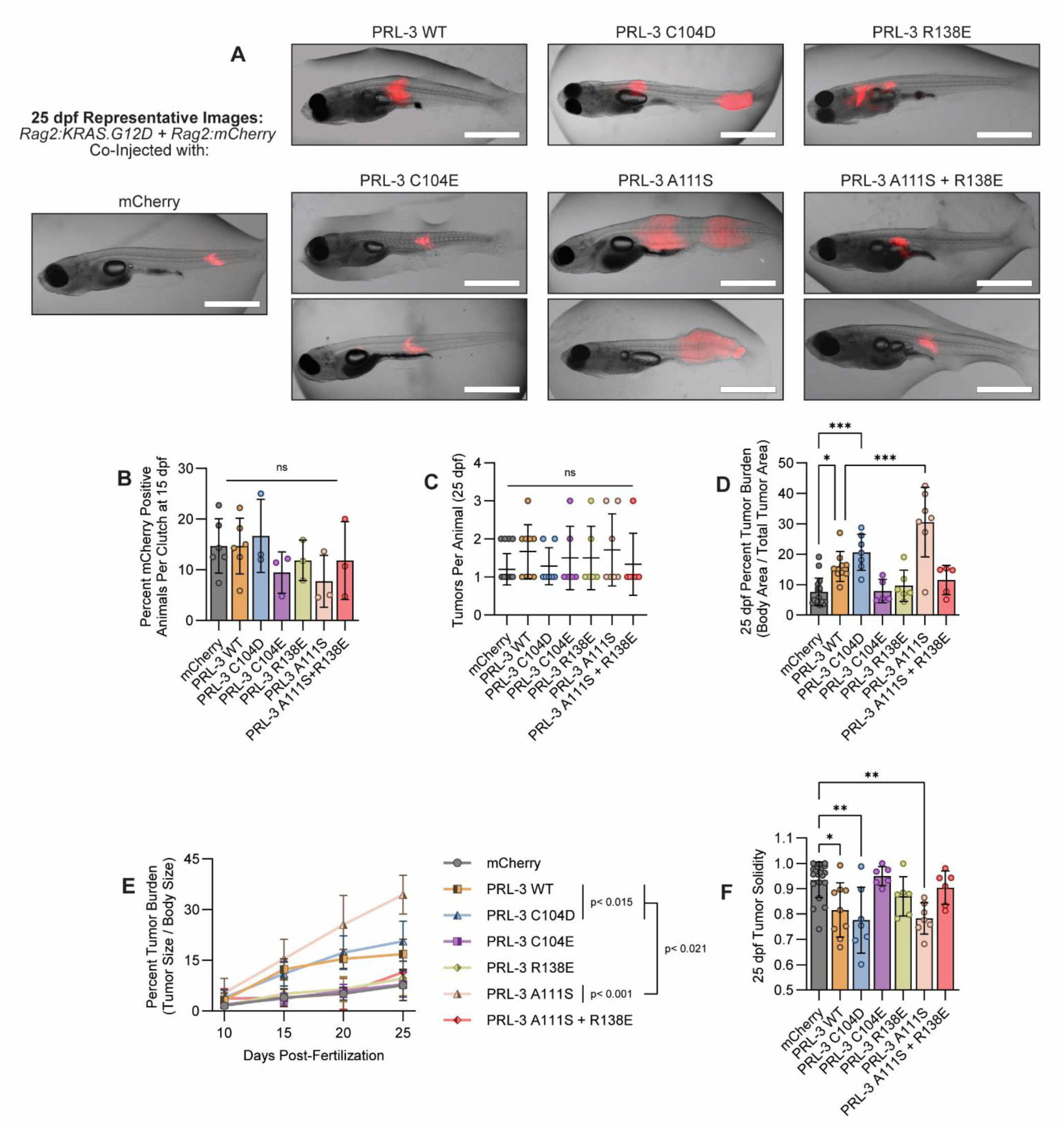
Enhancing the phosphatase activity of PRL-3 boosts tumor size in the RMS model in a CNNM-binding-dependent manner. **(A)** Representative images of animals injected with the indicated RMS constructs and PRL-3 mutants. **(B)** Quantification of animals per clutch showing detectable mCherry signal at 15 days post-fertilization (dpf); each data point represents one clutch (≥25 animals). **(C)** Quantification of the number of tumors per animal at 25 dpf. Animals with tumor burdens exceeding 50% of their total body size were excluded from analysis. **(D)** Quantification of tumor burden relative to animal body size at 25 dpf. Only the largest tumor was measured in animals with multiple tumors. **(E)** Quantification of tumor burden relative to body size over time. **(F)** Quantification of tumor solidity at 25 dpf. Tumors not located along the animal’s torso or tail (such as those on the dorsal fin or jaw) were excluded from the analysis due to differing morphology. Data in C-F were pooled from at least 3 clutches per group; each data point represents an individual animal. Error bars represent standard deviation. Statistical significance was assessed with an ordinary one-way ANOVA with Tukey correction (A, C, E-G), unpaired t-test (B), or two-way ANOVA (D). *, *p*<0.05; **, *p*<0.01; ***, *p*<0.001; ****, *p*<0.0001.

**Supplemental Figure 9.**
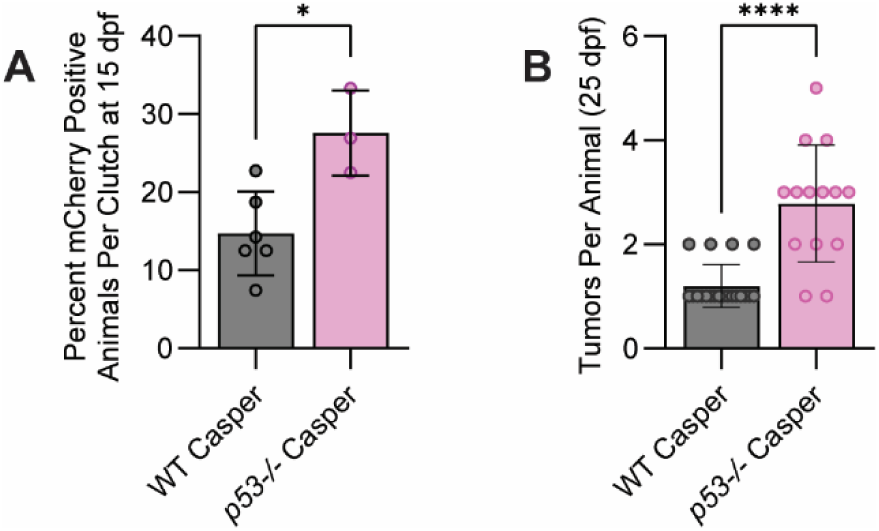
Knockout of p53 increases tumorigenesis in the RMS model. **(A)** Quantification of animals per clutch with detectable mCherry signal at 15 dpf, indicating tumor presence. Each data point represents an individual clutch of at least 25 animals. **(B)** Quantification of tumor number per animal at 25 dpf. Error bars represent standard deviation. Statistical significance was determined using an unpaired *t*-test. *, *p*<0.05; ****, *p*< 0.0001.

**Supplemental Figure 10.**
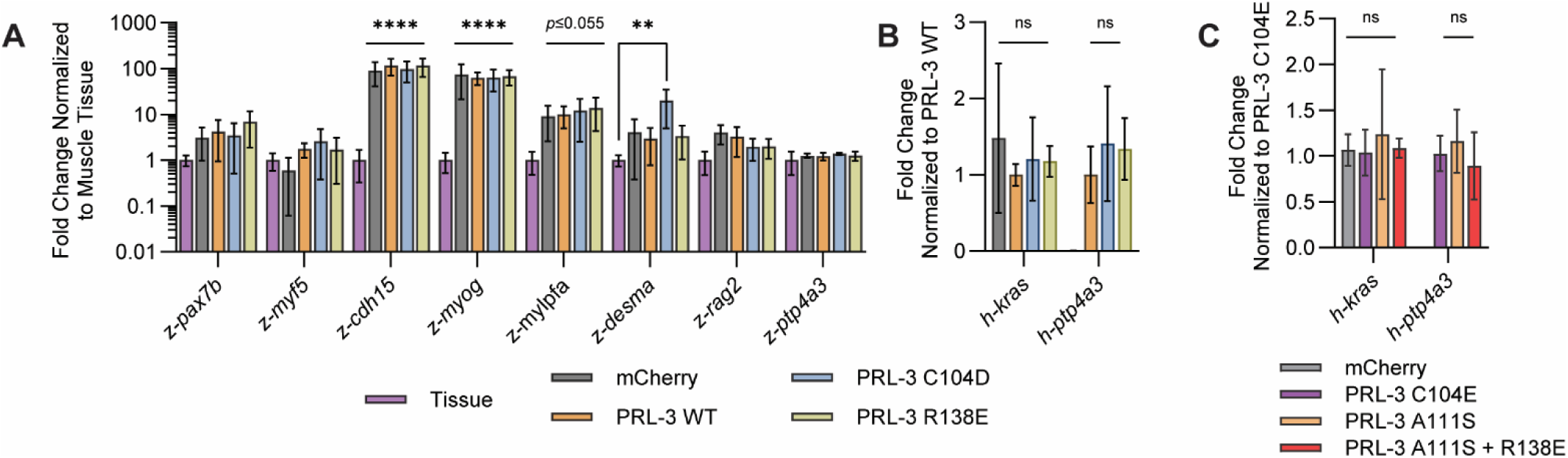
The overexpression of PRL-3 mutants has a minimal effect on RMS gene signatures. RT-qPCR results for endogenous (**A**) and injected transgenes (**B**) of interest in RMS samples, with expression normalized to non-cancerous muscle tissue. Each analysis was conducted with samples extracted from six animals per condition. (**C**) RT-qPCR analysis for injected transgenes, normalized to the PRL-3 C104E group, with three animals analyzed per condition. Error bars represent standard deviation. Statistical significance was determined by ordinary one-way ANOVA for each gene of interest, significance is defined as **, *p*<0.01; ****, *p*<0.0001

**Supplemental Figure 11.**
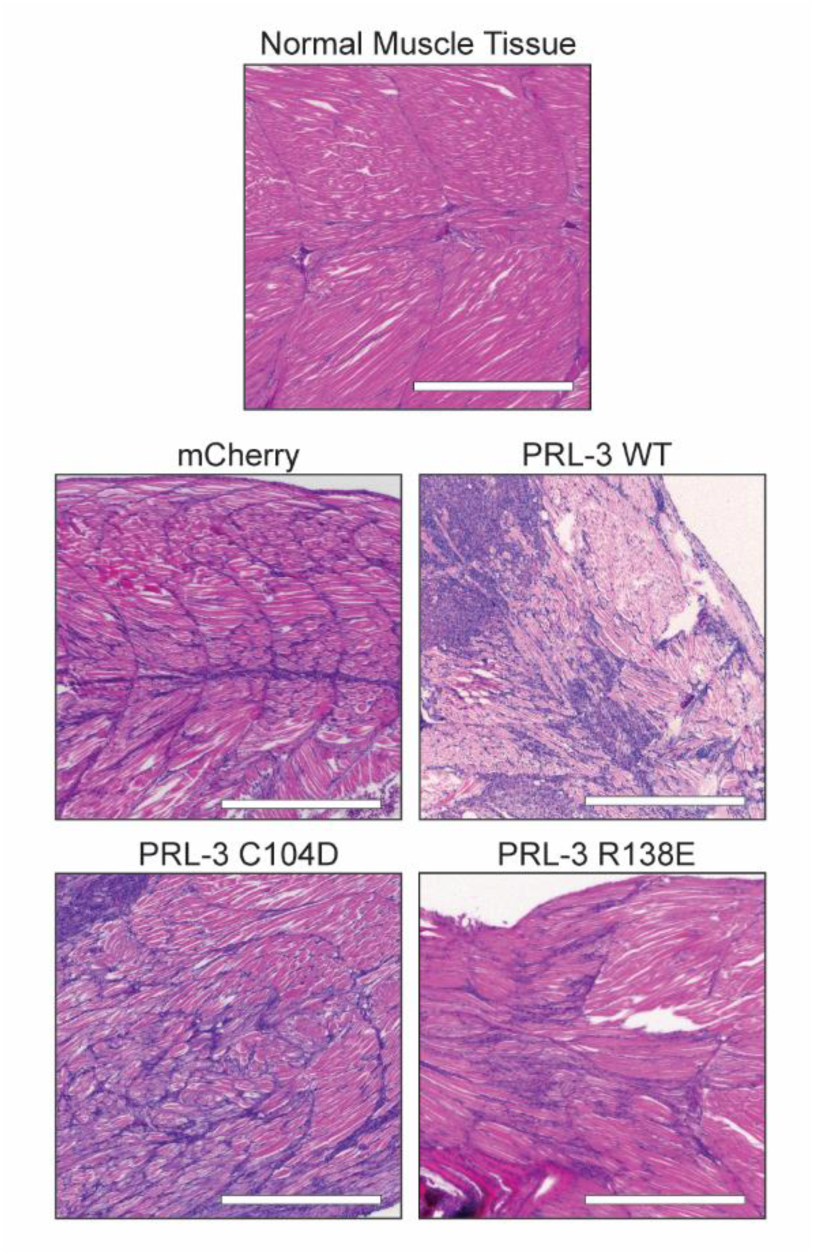
Representative H&E images of RMS tumor sections. Hematoxylin and eosin (H&E) staining of RMS tumor sections from zebrafish injected with rag2:KRAS(G12D) and the indicated PRL-3 transgene, shown alongside normal muscle tissue for comparison. Hematoxylin stains nuclei, while eosin stains cytoplasm and extracellular structures. Scale bar, 500 µm.

**Supplemental Figure 12.**
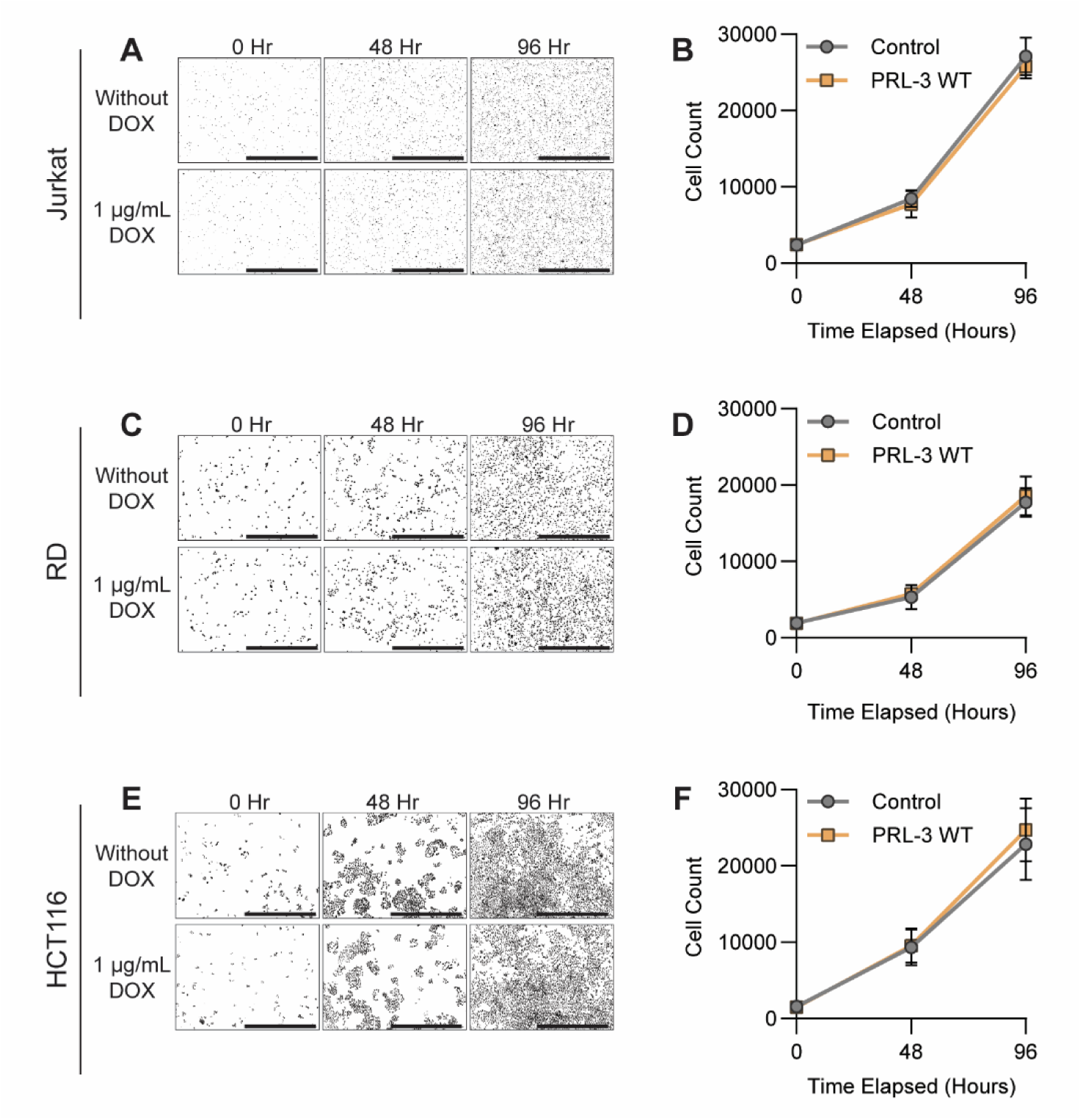
The overexpression of PRL-3 does not enhance proliferation in vitro. Representative images showing nuclei counts used to assess proliferation rates in Jurkat (**A**), RD (**C**), and HCT116 (**E**) cell lines with or without PRL-3 overexpression. **(B, D, F)** Quantification of cell numbers over time in the same cell lines under the same conditions. Data represent pooled results from at least three independent experiments for each cell line. Error bars indicate standard deviation. No statistically significant differences were observed between conditions, as determined by *t*-test.

**Supplemental Figure 13.**
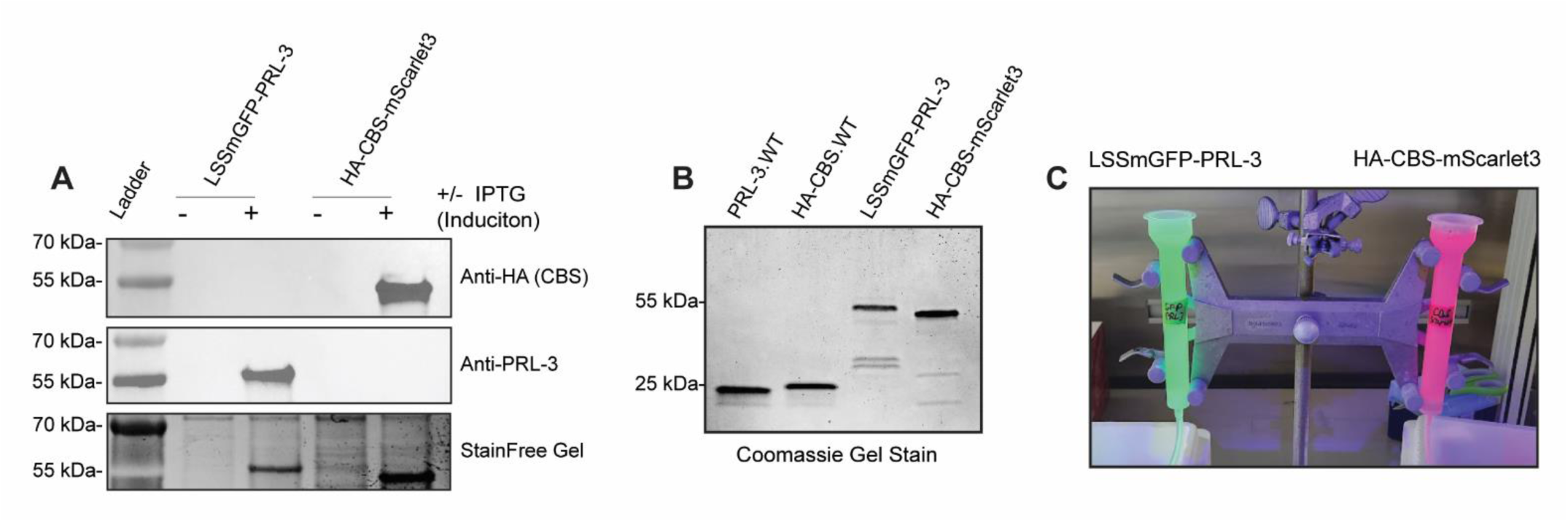
Purification of PRL:CBS FRET proteins. **(A)** Western blot showing recombinant protein expression following IPTG induction. **(B)** SDS-PAGE and Coomassie staining of purified proteins to confirm purity. **(C)** Fluorescence verification of FRET constructs (LSSmGFP-PRL-3 and HA-CBS-mScarlet) during the purification process.

**Supplemental Figure 14.**
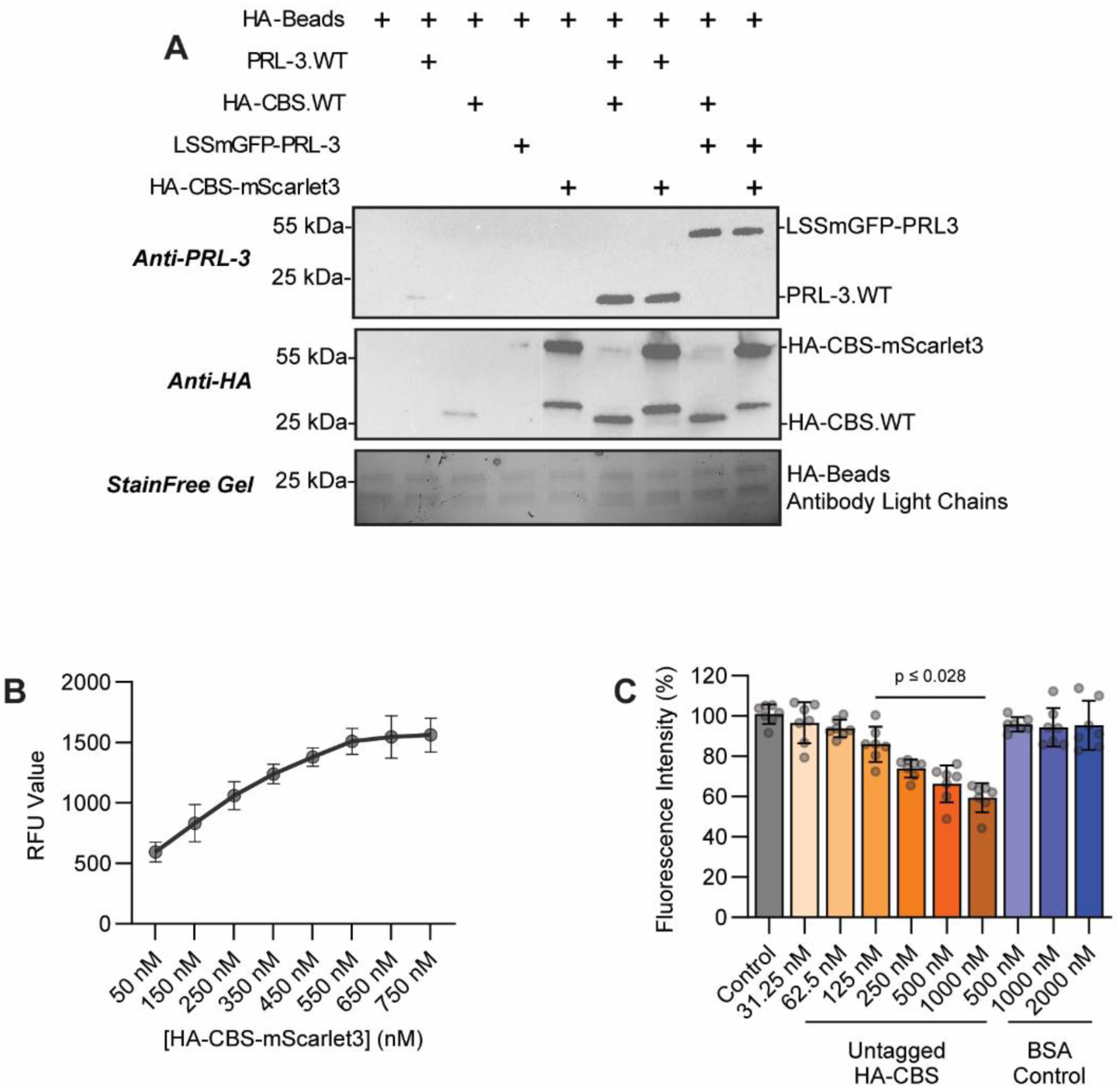
Validation of the PRL:CBS FRET pair. **(A)** In vitro immunoprecipitation assay using the FRET fusion proteins compared to their non-fluorescent protein counterparts. **(B)** FRET signal saturation observed upon addition of increasing concentrations of HA-CBS-mScarlet to a constant 250 nM concentration of LSSmGFP-PRL3. **(C)** Competitive FRET assay showing reduced signal with increasing concentrations of HA-CBS designed to compete for LSSmGFP-PRL3 binding and displacing HA-CBS-mScarlet. BSA was used as a negative control. Data are representative of two independent experiments, normalized to untreated controls. Error bars indicate standard deviation. Statistical significance was determined with ordinary one-way ANOVA and Tukey correction and is indicated in panel C.

**Supplemental Figure 15.**
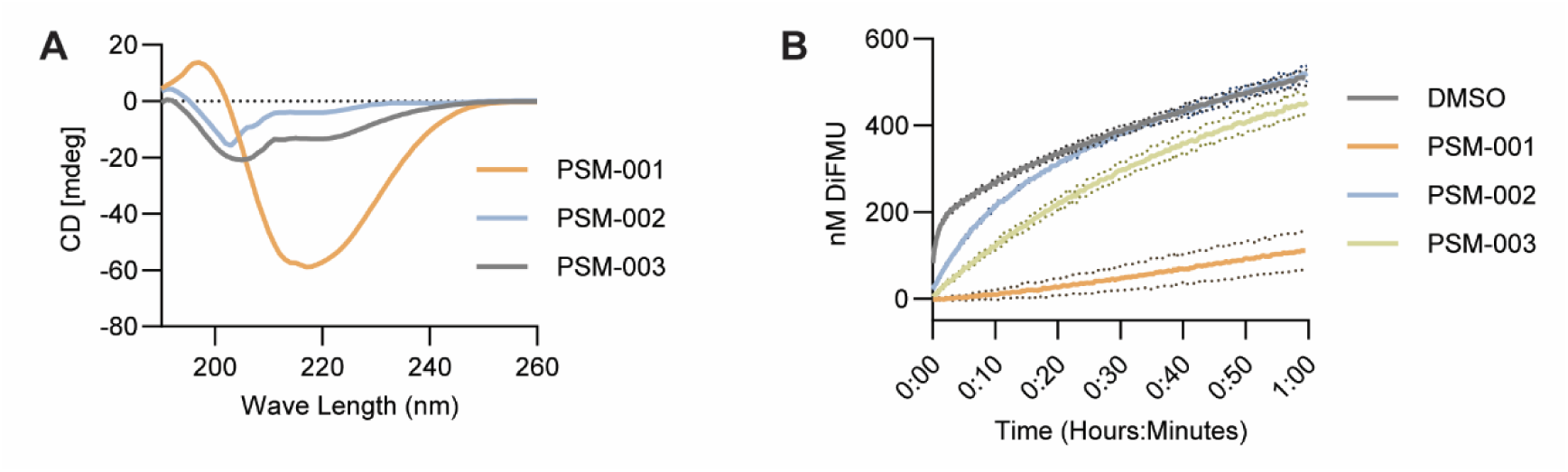
Peptide circularization strategies produce PRL-substrate mimics (PSMs) with variable structural and functional properties. **(A)** Circular dichroism (CD) spectra of each PSM peptide, illustrating differences in secondary structure. **(B)** DiFMUP phosphatase assay results assessing the inhibitory efficacy of each peptide against PRL enzymatic activity. The data shown are representative of at least two independent experiments. Dotted lines (B) indicate standard deviation.

**Supplemental Table 1:**
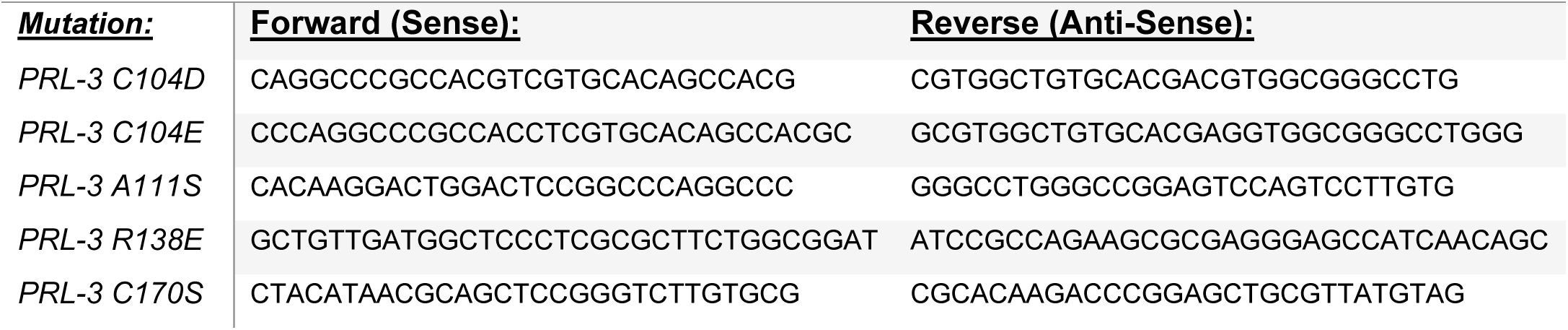
Site-Directed Mutagenesis Primers.

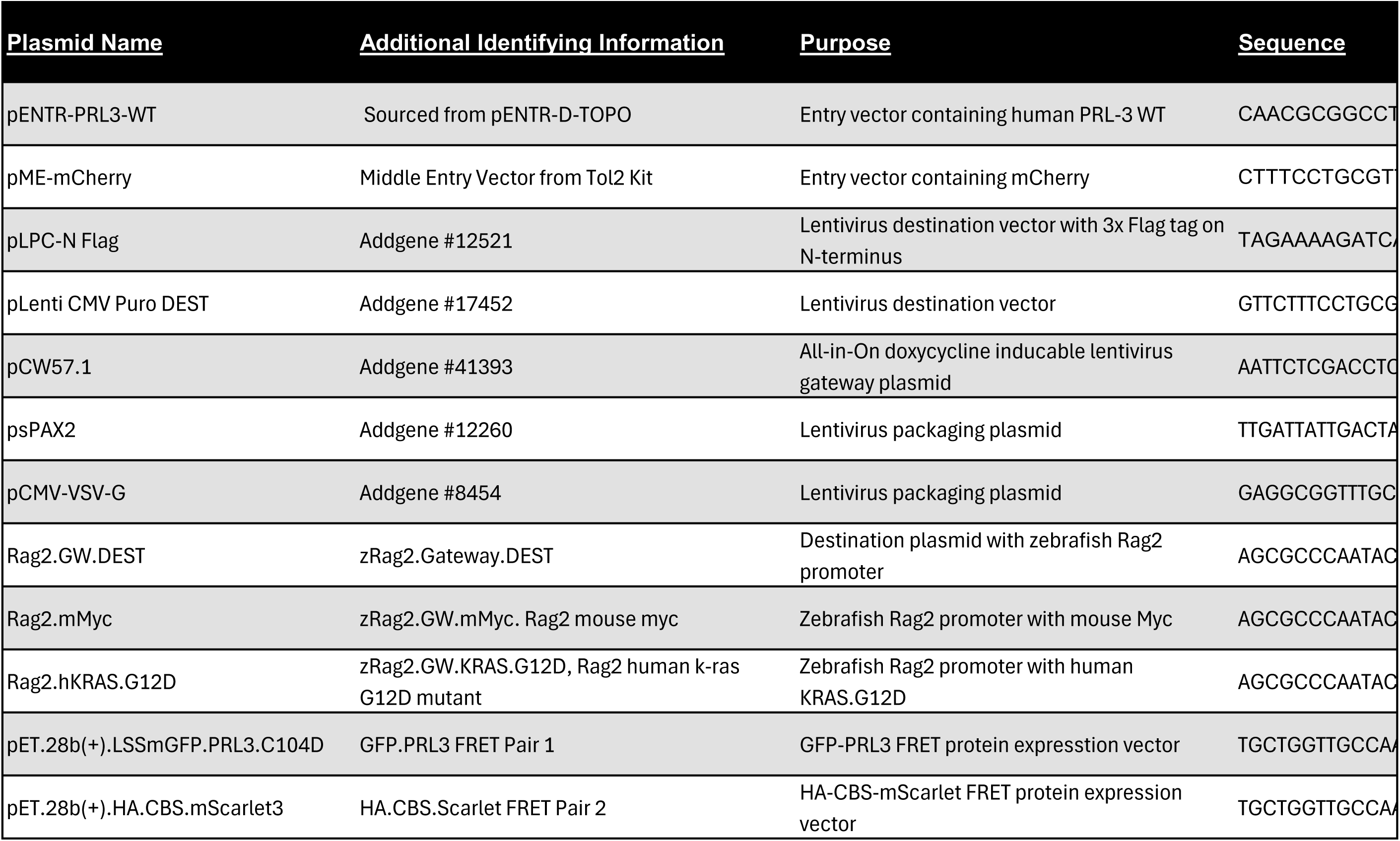

**Supplemental Table 3:**
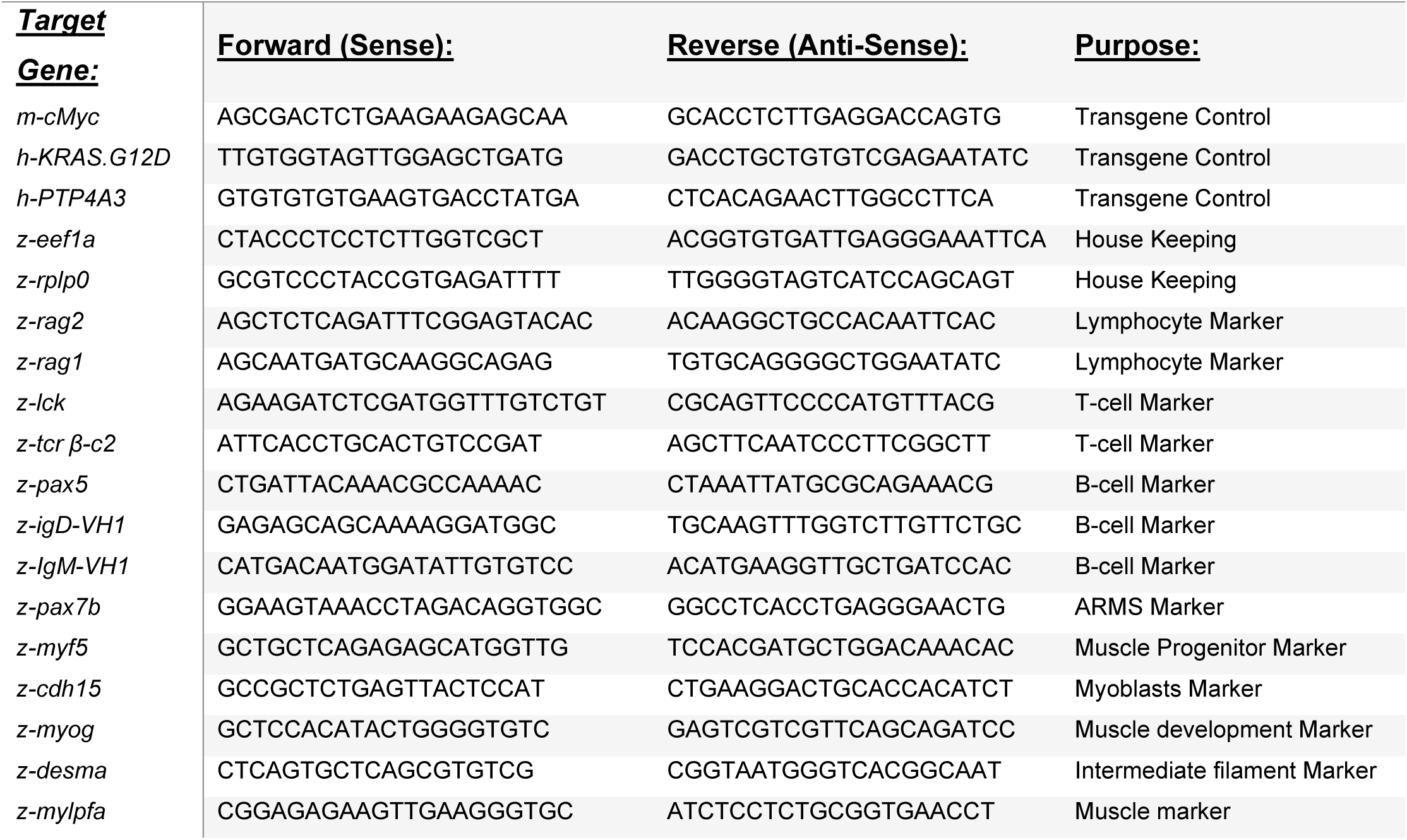
RT-qPCR Primers.

**Supplemental Table 4:**
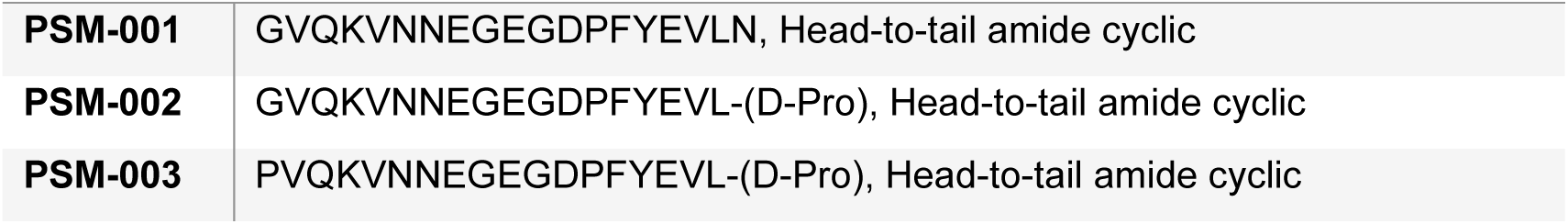
Cyclic Peptide Sequences and Linkers.

## Notes

### Competing Interest Statement

The authors have declared no competing interest.

